# Fighting or Embracing Multiplicity in Neuroimaging? Neighborhood Leverage versus Global Calibration

**DOI:** 10.1101/706747

**Authors:** Gang Chen, Paul A. Taylor, Robert W. Cox, Luiz Pessoa

## Abstract

Neuroimaging faces the daunting challenge of multiple testing – an instance of *multiplicity* – that is associated with two other issues to some extent: low inference efficiency and poor reproducibility. Typically, the same statistical model is applied to each spatial unit independently in the approach of massively univariate modeling. In dealing with multiplicity, the general strategy employed in the field is the same regardless of the specifics: trust the local “unbiased” effect estimates while adjusting the extent of statistical evidence at the global level. However, in this approach, modeling efficiency is compromised because each spatial unit (e.g., voxel, region, matrix element) is treated as an isolated and independent entity during massively univariate modeling. In addition, the required step of multiple testing “correction” by taking into consideration spatial relatedness, or *neighborhood leverage*, can only partly recoup statistical efficiency, resulting in potentially excessive penalization as well as arbitrariness due to thresholding procedures. Moreover, the assigned statistical evidence at the global level heavily relies on the data space (whole brain or a small volume). The present paper reviews how Stein’s paradox (1956) motivates a Bayesian multilevel (BML) approach that, rather than fighting multiplicity, embraces it to our advantage through a *global calibration* process among spatial units. Global calibration is accomplished via a Gaussian distribution for the cross-region effects whose properties are not a priori specified, but a posteriori determined by the data at hand through the BML model. Our framework therefore incorporates multiplicity as integral to the modeling structure, not a separate correction step. By turning multiplicity into a strength, we aim to achieve five goals: 1) improve model efficiency with higher predictive accuracy, 2) control the errors of incorrect magnitude and incorrect sign, 3) validate each model relative to competing candidates, 4) reduce the reliance and sensitivity on the choice of data space, and 5) encourage full results reporting. Our modeling proposal reverberates with recent proposals to eliminate the dichotomization of statistical evidence (“significant” vs. “non-significant”), to improve the interpretability of study findings, as well as to promote reporting the full gamut of results (not only “significant” ones), thereby enhancing research transparency and reproducibility.

## Introduction

Neuroimaging techniques often measure multiple signals simultaneously, such as from electro- or magneto-encephalogram sensors, or at spatial locations across the brain in functional magnetic resonance imaging (FMRI). In the former case, one may have tens to hundreds of nodes, while in the latter, one may have signals at over 100,000 spatial units (voxels and/or grayordinates) in the brain. A researcher will then be interested in evaluating how signals vary according to experimental conditions. This presents a particular kind of statistical problem, with fundamental issues that are still debated in the field after decades of existence.

In FMRI, the conventional analysis at the group level adopts a so-called *massively univariate analysis* approach with a two-step procedure^1^. First, one performs a statistical analysis at every location separately but usually with the same model (or design matrix). The analysis itself can be a simple *t*-test (do signals during conditions A and B differ convincingly?) or a more elaborate linear model (ANOVA, GLM, or mixed-effects model) with several predictors and interaction terms (is there enough evidence for a three-way interaction?). But testing at multiple locations simultaneously forces the investigator to rethink her approach of controlling for the overall false positive rate (FPR) under the null hypothesis significance testing (NHST) framework. One is not simply worried about errors at a single location but at all locations of the dataset *at the same time* – hence the need for a second step. This multiple testing problem has led to beautiful and creative solutions by the neuroimaging community in the past quarter of century, including random field theory (Worsley et al., 1992), Monte Carlo simulations (Forman et al., 1995), and permutation testing (Nichols and Holmes, 2001; Smith and Nichols, 2009). However, we believe that the current approaches have an important shortcoming: excessive penalties from correction for multiplicity due to modeling inefficiency.

Consider the two-step procedure. In the first step, by performing analysis at every location separately, one is assuming that all spatial units are unrelated to each other; in other words, they do not share information. Although the local relatedness (i.e., smoothness) among neighboring spatial units can be approximately accounted for during the second step of multiple testing correction, any similarity among brain regions is fully ignored. For example, from the Bayesian perspective neglecting such common information is equivalent to assuming that the effect can take values uniformly from −∞ to +∞. Instead, in the present framework a Gaussian distribution (whose parameters are determined by the data) of effect magnitudes is assumed. It is also worth noting that various approaches have been developed to take into consideration the spatial (and temporal) relatedness for localized statistical inference (e.g., Lindquist 2008; Bowman et. al, 2008; Derado et. al, 2010; Kang et al, 2012; Zhang et al, 2015). Nevertheless, a multiplicity issue exists because there are as many models specified as the number of the spatial units, leading to potential overfitting and model inefficiency. A model specifies how the data are related to the effect in question, such as the difference of two means in a *t*-test or GLM. Whatever type of model is adopted in the first step, the investigator is forced by the presence of simultaneous inferences to follow up with a second step to adequately control for the overall chance of misidentification in the reported results. As mentioned, a series of correction methods have been developed in the past.

In the present paper, we argue that such a massively univariate approach leads to information waste and inefficient modeling. There may be a misperception outside the neuroimaging community that statistical analysis with brain data is untrustworthy, as if multiple testing were usually not corrected due to the misreading of the oft-noted dead salmon study that won the Ig Nobel prize (Bennet et al., 2010). The reality might be the opposite because of the modeling inefficiency. The multiplicity challenges are far from trivial and remain active research topics, as evinced in a recent paper that stirred substantial controversy (Eklund et al., 2016). Against this backdrop, we propose an *integrative approach* that incorporates all the spatial units into a single, unified Bayesian multilevel (BML) model, leading to potentially improved inference efficiency through *global calibration*. The BML framework also facilitates transparency, reporting a study’s full results instead of artificial dichotomization (“significant vs. non-significant”), and model validation. Accordingly, the modeling framework resonates with critical assessments of the NHST framework which have intensified in recent years (Nuzzo, 2014; Wasserstein and Lazar, 2016; McShane et al., 2018; Amrhein et al., 2019).

## Controlling overall FPR under the conventional framework

We start with a brief review of how multiplicity is handled under NHST in neuroimaging. The most basic correction for multiple testing is obtained via Bonferroni correction, namely by raising the bar for statistical evidence at each spatial unit by dividing the desired significance level by the number of spatial units (e.g., setting *α* = 0.0000005 if tests will be performed at 100, 000 locations). However, this approach assumes that effects at every spatial location are fully independent, an assumption that is violated in the case of FMRI data, making this classical approach highly conservative in general (i.e., there are fewer independent tests than the number of voxels). Investigators recognized early on that the probability that a clique of spatial units (such as a cluster of voxels) would be “active” together was much smaller than that of individual units. Accordingly, such spatial relatedness (i.e., voxel adjacency) has motivated most methodologies aimed at addressing multiplicity. Typical correction efforts belong to two main categories: 1) controlling for family-wise error, so that the overall family-wise error(FWE) at the whole-brain or cluster level is approximately at the nominal value, and 2) controlling for false discovery rate (FDR), which targets the expected proportion of identified items (“discoveries”) that are incorrectly labeled (Benjamini and Hochberg, 1995). The two approaches are conceptually different in adjusting the overall statistical evidence. FDR can be used to handle a needle-in-haystack problem, where a small number of effects exists among a sea of zero effects in, for example, bioinformatics. Although considered more powerful in general at the cost of increased FPR (Shaffer, 1995), FDR is usually more conservative for typical neuroimaging data most likely due to the complication of spatial correlation, evidenced by its rare adoption in the literature compared to the FWE correction methods currently implemented in the field.

Multiple testing correction methods tend to be “add on” procedures that are separate from the individual statistical models of interest. Therefore, they contain an element of *arbitrariness*, as evidenced by the existence of varied “correction” techniques across the field. Conceptually, the situation is similar to that of defining an island using “pixels” in a satellite image as a piece of sub-continental land above a fluctuating sea level (Fig. 1): the definition of the island will depend on the water-level threshold adopted, which itself may vary with tide, season, geological time, and a minimal area setting (e.g., 10,000 square meters). In the conventional statistics framework, the thresholding bar ideally plays the role of winnowing the wheat (true effect^2^) from the chaff (random noise), and a *p*-value of 0.05 is commonly adopted as a benchmark for comfort in most fields and an imprimatur during the publication review process. However, a problem facing the correction methods for multiple testing concerns “arbitrariness”, which can be examined from at least four different perspectives. First, the threshold or the cut-off value itself is arbitrary; why not use 0.04 or 0.06, for instance? Thresholding is a binary operator that splits the outcome into categories in a way that in practice entails the belief that the effect is artificially divided into “real vs. non-existing”; that is, data are separated into those shown with “strong evidence” and “the rest,” which are hidden from the publications. Second, although particular correction methods are developed rigorously, differences in their respective assumptions may lead to different results – random field theory, cluster-based simulations and permutations have reasonable foundations for addressing multiple testing, but they have slightly differing results, in general. Third, the statistical evidence resulting from multiple testing correction will heavily depend on the analyst’s focus, or data space: whole brain, gray matter, a subnetwork, or a list of regions. Lastly, even though spatial relatedness is largely accounted for locally among neighboring voxels in all correction methods as noted above, the common information globally shared across the brain is fully ignored.

**Figure 1:**
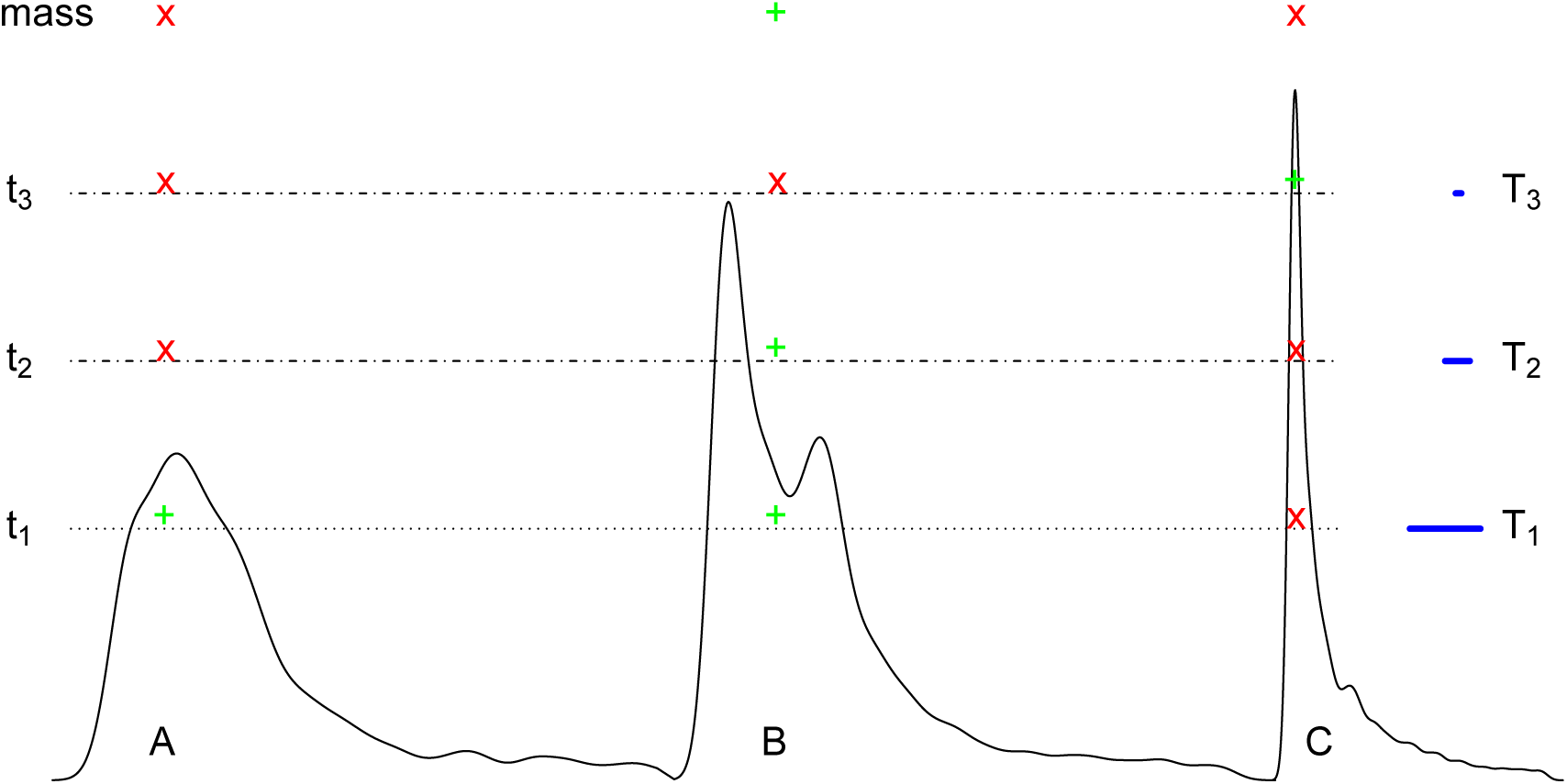
Schematic illustration of handling multiplicity via neighborhood leveraging procedures. The process is illustrated by considering three hypothetical spatial clusters that can be imagined as three sub-continental landmasses. The “pass” or “fail” of a cluster depends on its size (cross-section extent) at a particular voxel-wise threshold (or sea level; showcased by the blue segment length). In other words, only the neighborhood defined by that particular threshold matters in the sense that the spatiotemporal information contained in the data is summarized by a snapshot (a cluster defined by a cutoff). At the level of t1, clusters A and B survive (green “+” sign) based on the spatial threshold T1 while C does not (red “×” sign) because of its small size. In contrast, at the stringent level t3, both clusters A and B fail while C survives. Note that the surviving clusters strongly depend on the “sea level” adopted, and no single case is ideal. In particular, if an anatomical region is intrinsically small, clustering will often fail to reveal it unless the statistical evidence is unusually strong.

### Available methods

At present, there are four general approaches of neighborhood leverage to handling multiplicity in neuroimaging: two of them can be called “cluster-based” and two can be called “permutation-based”. The two cluster-based methods are Random Field Theory (as used in SPM^3^) and Monte Carlo simulations (as adopted in AFNI (Cox, 1996)): starting with a voxel-wise probability threshold (e.g., 0.01, 0.001) at the voxel level, a spatial-extent threshold is subsequently determined that specifies the minimal cluster size that should be believed, therefore controlling the overall FPR at the cluster level – the idea is that one should believe in the islands, not in the image pixels. The voxel-level *p*-value and the cluster size trade-off against each other, with a higher statistical voxel-level threshold leading to a smaller cluster size cutoff. Thus, a small region of activation can only gain ground with a low *p*-value at the voxel level, while large regions with a relatively large *p*-value at the voxel level may fail to survive the criterion (Fig. 1 at the threshold t_3_). Similarly, a lower statistical threshold (higher *p*-value) requires a larger cluster volume, so that smaller regions have little chance of reaching the survival level (Fig. 1 at the level t_1_ or t_2_). (For the investigator attracted to small regions of the brain, such as the amygdala or thalamic subnuclei, this poses a considerable headache.) The arbitrariness of the statistical threshold at the voxel level poses another challenge to the investigator: one typically will lose spatial specificity with a low threshold since small regions that are contiguous will become part of large clusters of activation (at times spanning hundreds of cubic milliliters in the brain); but note that the ability to detect large regions of activation is compromised when a high statistical threshold is chosen. A recent critique of the approach of cluster formation (Eklund et al., 2016) has resulted in a trend to require a higher statistical bar in neuroimaging (a voxel-wise threshold below 0.001). However, shifting the threshold does not affect the arbitrariness associated with the thresholding procedure itself: it is a corollary of dichotomization.

Alternative methods exist that employ permutation testing procedures. The maximum statistic approach (Nichols and Holmes, 2001), an early version, starts with the construction of a permutation-based null distribution of a maximum statistic (either maximum testing statistic or maximum cluster size) at a predetermined primary threshold. The original data are assessed against the null distribution, and the top winners at a designated rate (e.g., 5%) are declared as the surviving ones. While the approach is effective at maintaining the nominal FPR level, as in the parametric case above, the results will depend on the threshold employed, often strongly so. The maximum statistic approach has been extended to the analysis of matrix data, implemented in packages such as network-based statistics (NBS) (Zalesky et al., 2010) and the CONN toolbox (Whitfield-Gabrieli and Nieto-Castanon, 2012). Again, thresholding decisions may lead to disparate results when a different primary threshold is adopted – even as the same data are analyzed.

More recent permutation-based approaches take into consideration both effect magnitude and spatial extent (Smith and Nichols, 2009), and have been implemented in programs such as Randomise and PALM in FSL^4^ using threshold-free cluster enhancement (Smith and Nichols, 2009), and in 3dttest++ in AFNI using equitable thresholding and clusterization (Cox, 2019). A similar approach has also been borrowed in the analysis of matrix data (Bagio et al., 2018). These techniques circumvent the problem of having to choose a primary voxel-level threshold, thus ameliorating the issue of requiring a predetermined threshold in cluster-based methods. Nevertheless, the approach may come with some loss in statistical efficiency. For example, it is possible for a spatial clique to survive a cluster-based method that utilizes a primary threshold but not survive the permutation approach that combines both signal strength and statistical evidence (e.g., permutation versus cluster-based approach adopting a primary threshold at t_3_ in Fig. 1), especially when the spatial clique sizes are relatively homogeneous. On the other hand, the permutation approach might achieve higher sensitivity when the spatial size varies substantially (e.g., permutation versus cluster-based approach adopting a primary threshold at t_2_ in Fig. 1).

Regardless of the specifics of each adjustment method, the effect of interest is assessed through a model applied separately at each spatial unit under the current modeling framework. Yet, each individual unit is only meaningful when it belongs to a surviving cluster. Accordingly, the investigator cannot attach a specific uncertainty level (e.g., confidence interval under NHST) at particular locations because of the indivisible nature of the spatial cluster (Woo and Wager, 2015). As what survives is the entire cluster, the statistical significance at the voxel level is undefined.

### Other issues

Another challenge facing investigators involves potential “mid-analysis” changes of correction method, or more broadly of “data domain”. When a whole-brain voxel-wise analysis fails to allow a region to pass the peer-accepted threshold, is it now legitimate to go through additional analysis steps? For example, the existing literature may legitimately point towards a set of target brain regions, so should the investigator now focus on those regions, or possibly even a single “star” brain area (e.g., amygdala, accumbens, fusiform gyrus, etc.)? Changing the *model space*, such as the space of multiple testing, may seem to “improve” the strength of statistical evidence (Kruschke, 2010), despite the fact that the original data remains exactly the same! Whereas trying multiple analysis strategies until “statistical significance” is reached is clearly unacceptable, where should the line be drawn between legitimate “trying to understand one’s data” and walking in “the garden of forking paths”, “data snooping” or *p*-hacking? To claim that a study would come to a halt based on a “first pass” of data analysis is admittedly naive, and does not acknowledge the actual way that science proceeds in the laboratory.

A final issue with the massively univariate modeling strategy is that it implicitly makes the assumption that the investigator is fully ignorant about the distribution of effect sizes across space. Specifically, conventional statistical inferences under NHST assume that all potential effects have the same probability of being observed; it is assumed that they follow a uniform distribution from −∞ to +∞ (Gelman et al., 2014). However, in practice, many types of data tend to have a density of roughly Gaussian characteristics with a bell-shaped distribution exhibiting a single peak and near symmetry (which is not surprising given that the central limit theorem applies in many situations). The information waste resulting from the assumption of uniform instead of Gaussian distribution leads to inefficient modeling due to overfitting. The assumption of uniform distribution of effect sizes should be viewed in contrast to the adoption, indeed in the same conventional methods of statistical inference, of a Gaussian distribution when experimental entities are concerned (e.g., participants, animals). In fact, per the maximum entropy principle, the most conservative distribution is the Gaussian if the data have a finite variance (McElreath, 2016). Accordingly, the question that needs to be considered is as follows: Do we truly believe that effects across spatial units are uniformly dispersed across all possible values? If the answer is “no” (e.g., extremely high or low values are nearly unattainable), why cannot we assume that the effects across spatial units follow a Gaussian, as routinely practiced for the measuring units of subjects that are considered as random samples from a hypothetical pool, instead of adopting the stance of “full ignorance”? To anticipate, the Bayesian framework described here allows distributional assumptions to be incorporated in a principled manner, and to be validated against the observed data.

## Stein’s paradox and hierarchical modeling: two case studies

Charles Max Stein (1956) discovered an interesting but counterintuitive phenomenon in statistics and in decision theory: When three or more effects are estimated individually, the overall accuracy of each is worse than an integrative approach that shares information across the effects.

### Case study 1: predicting basketball player performance

Suppose that the shooting rate of Kevin Durant’s field goals during the 2018-2019 National Basketball Association (NBA) season is 52.1%. If one has to guess his field goal rate during the next season without any other information, his current efficiency would be the best (unbiased) estimate, even though his future performance may be better or worse, of course. The same would be true for any other player. Now consider the prediction of the future success rate of Durant as part of the top 50 NBA players. The surprising fact is that more accurate predictions can be produced compared to those simply using their current percentages as future predictions, as rigorously proven (James and Stein 1961; Efron and Morris, 1976). In other words, by considering each player as a member of a pool of the top 50 performers, we can make overall more accurate predictions than directly using each player’s current performance as a prediction (Fig. 2). Thus, although we can reasonably assume that all players’ performances are independent of one another, it is better to pool information across all players to make a prediction for each individual player. The pooling is considered “partial” when some information is borrowed from other players when predicting each individual player, in contrast to “complete” pooling^5^ in which all players are assigned the same prediction (e.g., overall average). The paradox was later substantially elaborated and rigorously extended by Efron (e.g., Efron and Morris, 1976). Although Stein’s paradox was originally proven for the case of Gaussian distributions, it has since been generalized to other distributions and even multivariate cases.

**Figure 2:**
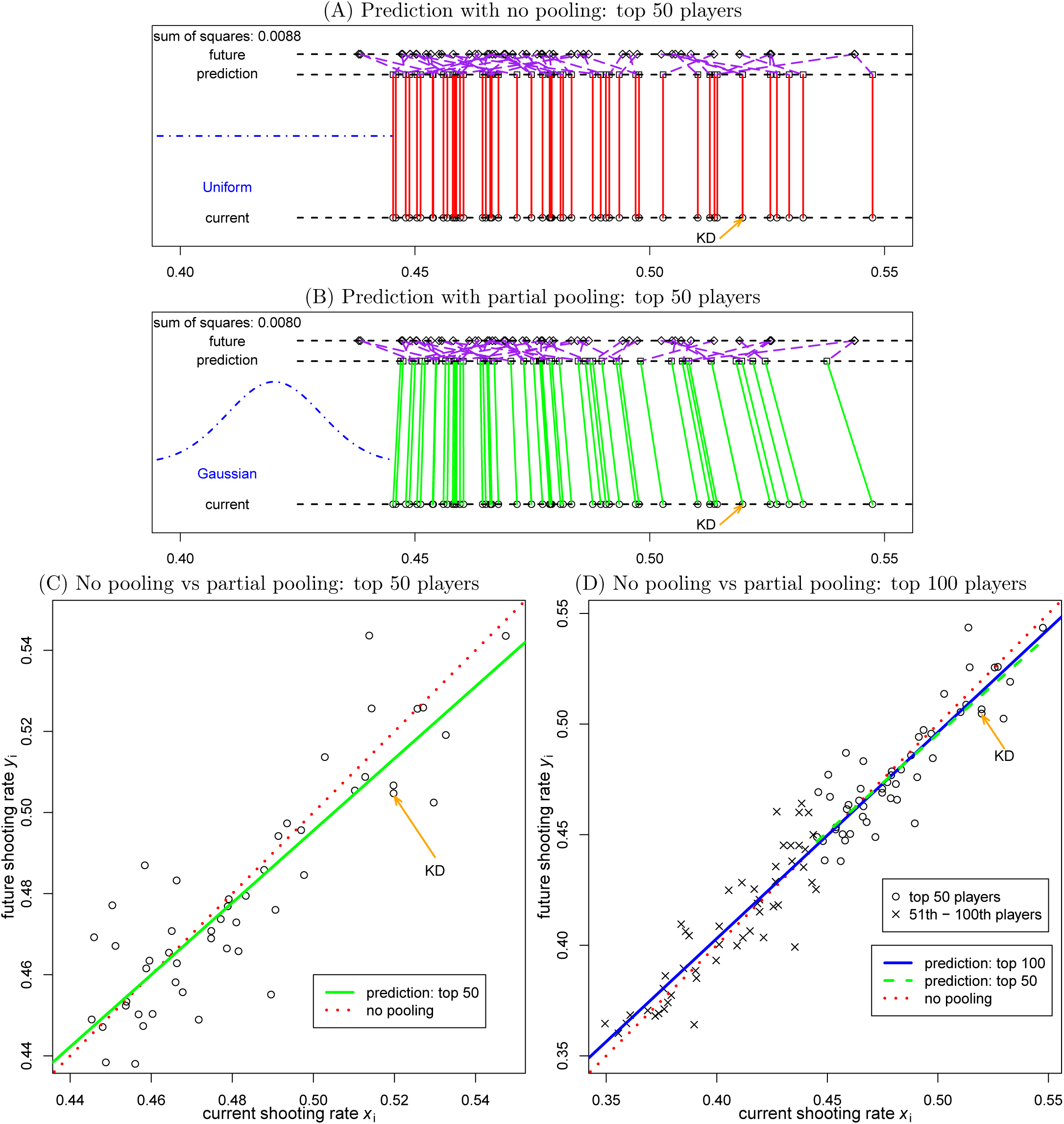
Geometric illustration of Stein’s paradox with simulated data of the top basketball players. Here, we pretend that we could time-travel to the future and take a sneak peek at the independent shooting rates *y*_*i*_ of the players, which are assumed to follow a Gaussian distribution: 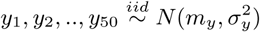 (here, *m*_*y*_ = 0.4, *σ*_*y*_ = 0.1). Kevin Durant’s data point is labeled as “LBJ”. (A) Simple scheme in which future performance of the top 50 players is predicted (values represented by squares) to be the same as in the current year (circles). The actual “future” values are also shown (diamonds). The purple dash lines link the predicted and actual future values. (B) The performance of the top 50 players is predicted through simple regression. The slanted green lines, corresponding to the green fitted line in (C), illustrate partial pooling, which interestingly also corresponds to the concept of regression to the mean established by Francis Galton. For the hypothetical dataset at hand, note that the prediction of the next season for Durant as one of the top 50 players is 49.0%, a value that is slightly downgraded from his current 52.1%. (C) Two scenarios are contrasted for the top 50 players. First, the same as in panel (A): using each player’s current performance to predict future values gives the diagonal line fit *x* = *y* (red line). Second, we could imagine having future data and predicting future *y*_*i*_ values with the current data *x*_*i*_ as an explanatory variable. In this case, the ordinary least squares solution (green line) would produce a fitted line shallower than the diagonal (dotted red line), which illustrates the partial pooling effect (see, Stigler 1990). (D) What would happen if we predicted the performance of the top 50 players as part of a bigger pool of the top 100 players? The effect of partial pooling remains evident by the shallower line of regression fit (sold blue line). Although the prediction of the top 50 players as part of the top 100 players is different from that limited to the top 50 players (dashed green line, same as the solid green line in panel C), the prediction difference is relatively small due to the adaptive nature of the Gaussian distribution assumed for the data.

The moral we can learn from the NBA player example is that the fundamental difference between the two modeling approaches lies in our willingness to apply available knowledge. The at-first intuitive approach of using each player’s current performance to predict future percentage (no pooling, Fig. 2A; note that current and predicted values are the same) assumes that we do not have any knowledge about the distribution of performances, and thus treat each player as an independent entity. In fact, such stance of “total ignorance” is equivalent to making the assumption that shooting rate performance is uniformly distributed. That is to say, all possible values are credible, including very low (e.g., 10%) and very high (e.g., 90%) shooting rates, which are actually not observed in practice.

It is instructive to consider the problem of simultaneously predicting the performance of the set of 50 players as a type of multiplicity problem. In this case, it is not only the individual accuracy (cf. voxel-wise *p*-value under NHST) that matters the most, but the overall accuracy (cf. overall FPR) that is of concern. As stated, one approach is to treat each player as independent from the rest, similar to the massively univariate approach in neuroimaging (Fig. 2A). As players do not simply repeat their performance, the observed future performance (generated hypothetically) will deviate from the preceding season to some extent (as indicated by the purple lines connecting squares to diamonds in Fig. 2A). We go through this exercise again but now pretend that we could time-travel to the future and take a sneak peek at the independent shooting rates *y*_*i*_ of the top 50 NBA players. If we predicted players’ future performance based on current performance via linear regression, we would obtain the fit displayed in Fig. 2C (green line). The same data as used in Fig. 2A were employed, which was generated based on the assumption that future shooting rates follow a Gaussian distribution. If we plot the linear regression predictions in a manner that follows that of Fig. 2A, we can see that the predicted values differ from the current one, as can be seen by the slanted green lines in Fig. 2B. The role of the Gaussian distribution assumption of future performance can be visualized as a sort of elastic band, whereby predictions are mutually informative. Notably, predictions for outlying players (high and low scorers) are shrunk toward the group mean via the implicit partial pooling of the procedure.

In the present case, our willingness to apply prior knowledge (that shooting percentages typically follow a Gaussian distribution) transformed the problem into a simple GLM system, thereby achieving a higher overall accuracy in Fig. 2B (see the comparison between the two sums of squared residuals) than that with the assumption of no prior knowledge (represented by a uniform distribution) in Fig. 2A. Notably, the prior information (i.e., the distribution assumption) only sets the general shape while the specifics of the shape (i.e., mean and variance) can be determined from actual data through the model (and estimated via ordinary least squares, or maximum likelihood). Thus, partial pooling is adaptive in the sense that information about centrality and spread of the players’ performances is not required as part of the prior information, but is determined from the data. The adaptivity of the Gaussian prior can be further illustrated as follows. The prediction for the top 50 players through regression is different when they are considered as a subset of the top 100 players. However, the difference due to the addition of the 51th-100th players is relatively small (Fig. 2D).

### Case study 2: dealing with epidemiological data of kidney cancer

An example conceptually closer to the situation in neuroimaging involves epidemiological survey data regarding the highest and lowest cancer rates across the United States (Fig. 3). Examination of the map indicates that the highest kidney cancer rates were observed in relatively sparsely populated regions of the country (Fig. 3A). One then might be tempted to infer that geographical factors explain the high rates, such as restricted access to healthcare, low utility infrastructure, or higher doses of radiation. But it is difficult to reconcile these explanations with the observation that counties with the lowest kidney cancer death rates also tend to be less populated (Fig. 3B). A more parsimonious explanation of the data, which is consistent with both maps, is based on the small sample size of rural counties, combined with the relatively low mortality rate due to kidney cancer (which is a rare disease). To see this, suppose that the national average death rate is around 1 per 2,000 people (per 10 years). In a county with a population of about 1,000 people, one can imagine observing zero or one deaths by chance alone, which would place the county into the map with the lowest or highest rate, respectively.

**Figure 3:**
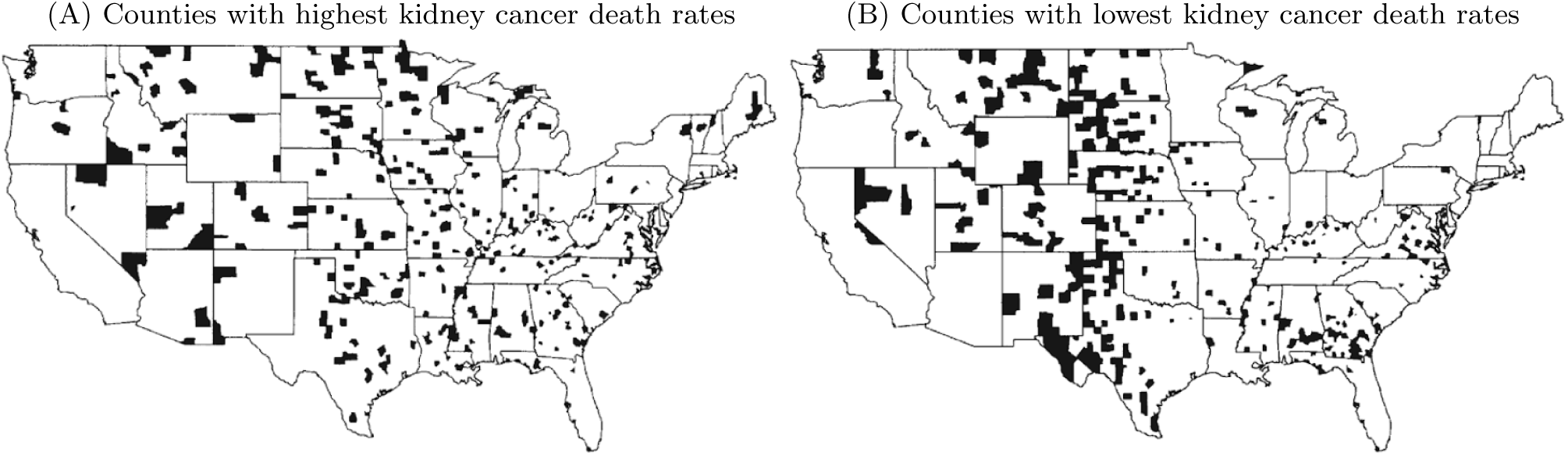
The U. S. counties with the highest (A) and the lowest (B) 10% death rates for kidney cancer during 1980-1989 are shown with blots. Republished with permission of Taylor & Francis Group LLC, from Gelman et al. (2013); permission conveyed through Copyright Clearance Center, Inc.

So, based on the survey data, how should we make reasonable inferences about *all* the individual counties? Two questions can be posed: 1) Should we really believe that a resident in a specific county has a higher (or lower) probability of dying of kidney cancer than the national average? 2) Can we infer that a resident in a county in Fig. 3A (or Fig. 3B) has a much higher (or lower) probability of dying of kidney cancer than a person in a county in Fig. 3B (or Fig. 3A)? That is, would we be able to confidently recommend that someone move from a county in Fig. 3A to one in Fig. 3B, or to discourage someone from moving from a county in Fig. 3B to one Fig. 3A? The answers would appear to be “no” in both cases.

The epidemiological distribution of kidney cancer deaths can be seen as involving a multiplicity problem over the several thousands of districts. A natural solution might be to borrow the idea of neighborhood leverage in neuroimaging (that is, spatial clustering), and to utilize the relatedness among adjacent counties to modulate the statistical evidence for clusters of counties. Note, however, that this leveraging approach only adjusts the extent of statistical evidence, and it does not modify the death rate estimates themselves (at county or cluster level). In other words, the conventional neuroimaging approach to handling multiplicity does not address two types of estimation errors that are important to consider (Gelman and Tuerlinckx, 2000). One is the so-called *type M* error, which refers to a potential over- or under-estimation of the effect magnitude. The second is the so-called *type S* error, which refers to getting the sign of a comparison wrong (does county A have a higher or lower death rate than county B?). A potential solution is to employ a methodology that prevents the estimation process from being swayed by larger fluctuations.

Consideration of Stein’s paradox in the context of the kidney cancer case suggests potential strategies to address the multiplicity issue. One possibility is to not place trust only in isolated bits of survey data (at the county level), but instead to incorporate all counties into a Bayesian model with a hierarchical structure that leverages death rate information across counties (Gelman et al., 2014). The BML model reflects the central concept behind Stein’s paradox: We should be somewhat sceptical about individual inferences of effect estimates and their uncertainties, especially extreme ones. Furthermore, we should adopt some information sharing – what is called partial pooling or shrinkage because of the tendency to reduce extreme values – which is possible through a hierarchical model that shares information across all, not just neighboring, spatial units. One may feel uneasy about partial pooling because of its introduction of *bias* to individual effect estimates^6^. The seemingly puzzling phenomenon of Stein’s paradox aptly captures the biased predictions about Durant’s future performance among his NBA peers (remember that the unbiased estimates should follow the *y* = *x* red dotted lines in Fig. 2). In the example, we shifted our focus from the accuracy of one particular effect (Durant’s future shooting rate; or, in the cancer case, the death rate at one specific county) to the *overall* predictive accuracy among all effects (overall accuracy among all the 50 players or among all US counties).

### BML: trading off bias against predictive accuracy

To recapitulate, a fundamental difference between the conventional, isolated modeling approach and the integrative BML method is that the former seeks statistical unbiasedness at each spatial unit, while accepting a daunting FPR problem that requires corrective procedures. Having an unbiased estimator (the quintessential example is probably the sample mean as an estimate of the population mean) may sound appealing, but it is not always the most desirable property, as recognized by statisticians and applied scientists for some time. This is especially the case when sample size is not large or when noise potentially overwhelms the effect, conditions all too often encountered by experimentalists in general and neuroimagers in particular. Briefly, in many circumstances, an unbiased estimator may have high variance. In such cases, it can fluctuate non-trivially from sample to sample. Adopting a biased estimator, instead, may be beneficial if it has lower variance; thus, although the estimator does not tend, in the limit (in practice with rather large sample sizes) to the “true” value of the effect of interest, it can prove advantageous in many practical settings. In fact, shrinkage estimation has been effectively applied to parcellating brain regions at the subject level by borrowing strength across the whole group of subjects (Mejia et al., 2015).

As schematically demonstrated in Fig. 2, the introduction of a Gaussian distribution introduces a mild constraint when estimating all players’ performances simultaneously (their shooting rates are not free to vary everywhere, but are informed by the entire set of performances in a Gaussian fashion). In fact, many conventional methodologies employ constraints to their estimation procedures. More technically, they benefit from regularization (Tikhonov et al., 1998), which involves introducing additional information in order to solve an ill-posed problem or to prevent overfitting. For example, ridge regression, LASSO (Tibshirani, 2009), elastic nets, and linear mixed-effects framework introduce additional constraints to the estimated regression coefficients in addition to minimizing residuals (e.g., the LASSO attempts to keep the overall “budget” of weights under a certain value: ∑_*j*_|*β*_*j*_| ≤ *k*). It is also interesting to note that global calibration is employed in meta analysis (Glass, 1976). When summarizing multiple studies of an effect, there is no correction for multiplicity of statistical evidence, but instead effects are weighted based on their respective relative reliability.

Some properties of the BML to be described further below include the following. First, the multiplicity issue is automatically dissolved given that a *single* model is employed – there is no “multiple” to correct! In addition to being conservative about extreme effects due to the shrinkage phenomenon, there is only one overall posterior that is formulated as a joint distribution in a high-dimensional parameter space (the dimensionality of which equals the number of parameters in the model). No multiplicity is incurred because 1) all the spatial units are included in one model and regularized through a reasonable prior, and 2) only one overall posterior distribution is utilized to predict various effects of interest. In fact, the analyst can make inference for as many effects of interest (including comparisons) as desired because each effect of interest is simply a marginal distribution (or a perspective projection) of the overall high-dimensional joint posterior distribution (like various angles of projecting the earth onto a 2D map). Second, an important focus becomes addressing type M and type S errors (see Fig. 4) (Gelman and Tuerlinckx, 2000), both of which are meaningful in practice. Third, model efficiency, misspecifications and performance can be directly assessed and compared straightforwardly through predictive accuracy and cross validations, similar to techniques such as least squares, maximum likelihood and information criterion. One predictive accuracy indicator is the Watanabe-Akaike information criterion, for instance, via leave-one-out cross-validation (Vehtari et al., 2017), and another one involves “posterior predictive checks”, which simulate replicated data under the fitted model so as to graphically compare actual data to model predictions.

**Figure 4:**
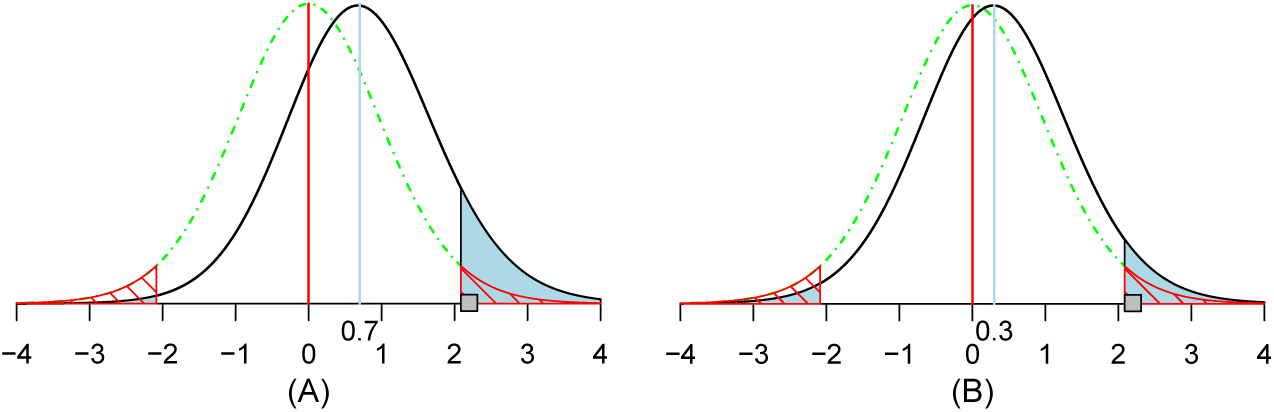
Illustration of the concept and interpretation for type I (FPR), type II, type S and type M errors. Suppose that there is a hypothetical noncentral *t*_20_ distribution (solid black curve) for a true effect (blue vertical line) of 0.7 (A) or 0.3 (B) and a standard error of 1.0. Under the null hypothesis (red vertical line and dot-dashed green curve), two-tailed testing with a type I error of 0.05 leads to having thresholds at ±2.09; FPR = 0.05 corresponds to the null distribution’s total area of the two tails (marked with red diagonal lines). The power (shaded in blue) is the total area of the *t*_20_ distribution for the true effect (black curve) beyond these thresholds, which is 0.10 (A) or 0.06 (B). The *type S* error is the ratio of the blue area in the true effect distribution’s left tail beyond the threshold of −2.09 to the total area of both tails, which is 4% (A) or 21% (B) (i.e., the ratio of the “statistically significant” area in the wrong-signed tail to that of the total “statistically significant” area). If a random draw from the *t*_20_ distribution under the true effect happens to be 2.2 (small gray square), it would be identified as statistically significant at the 0.05 level, and the resulting *type M* error would quantify the magnification of the estimated effect size as 2.2*/*0.7 ≈ 3.1 (A) or 2.2*/*0.3 ≈ 7.3 (B), which is substantially larger than unity. These two plots aptly demonstrate the importance of controlling the errors of incorrect sign and incorrect magnitude when a large amount of variability exists in the data. High variance is bound to occur when we deal with the multiplicity issue embedded with massively univariate modeling, which is further exacerbated by scenarios such as small sample size, noisy data, unaccounted for cross-subjects variability and suboptimal alignment to a standard template. More systematic exploration and comparison between the conventional (type I and type II) and the new (type S and type M) sets of errors can be found in Gelman and Tuerlinckx (2000) and Chen et al. (2019b).

## Focusing on the research hypothesis instead of fighting a “strawman”

Before applying the BML approach to FMRI data, it is instructive to briefly compare the frameworks of NHST inference (as commonly employed by experimentalists) and the proposed Bayesian approach. Consider a scenario in which a single one-sample *t*-test with 20 degrees of freedom (e.g., 21 subjects) is employed in the NHST setting. The null hypothesis *H*_0_ is that the population mean is zero. Suppose that the data indicate that *t*_20_ = 2.85. If *H*_0_ were true, the probability of observing a *t*-value so extreme is rather low (*p* = 0.01 in Fig. 5, left). The *t*-value value thus provides a measure of “surprise”: How surprising would it be to observe such an extreme value in a world in which *H*_0_ were really true? In this setting, one estimates the extent of surprise *P* (data | *H*_0_). By custom if *p <* 0.05, one declares that the effect is “present”, with the all-important “significance” label.

**Figure 5:**
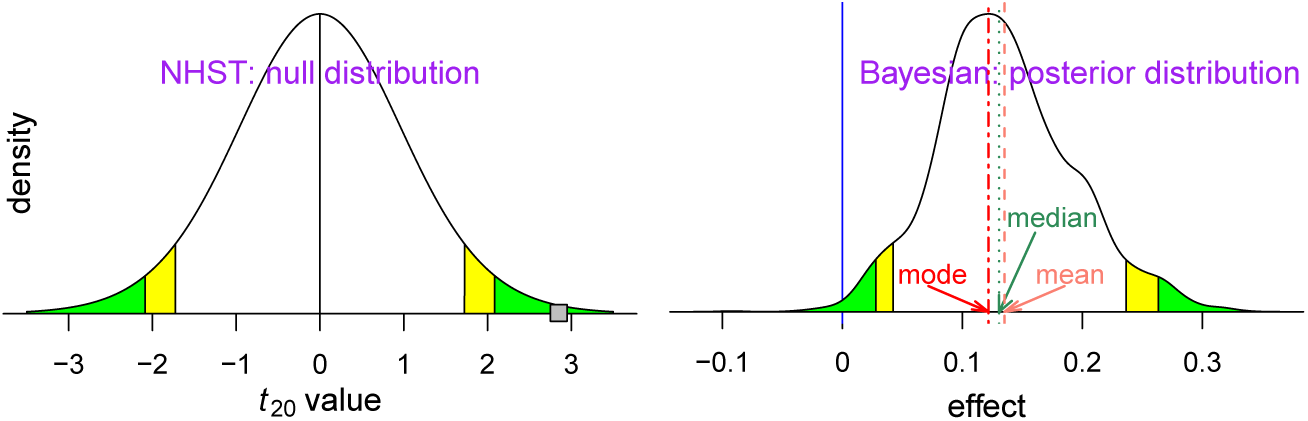
Different probability definition and focus between conventional and Bayesian frameworks. Statistical inferences under NHST usually pivot around the “weirdness indicator” of the *p*-value (left curve): green (or yellow) tails symbolize a two-sided significance level of 0.05 (or 0.1). If the data renders a *t*_20_ value of 2.85 (small gray square), reaching the comfort zone (green or yellow area) with a two-tailed *p*-value of 0.01 (the probability of obtaining such data with a *t*_20_ value at least as extreme if the effect were truly zero), one may declare to have strong evidence for the effect with an FPR threshold of 0.05 under the NHST framework. In contrast, inferences under the Bayesian framework directly address the research interest (right curve): what is the probability of the effect magnitude being greater than 0 with the data at hand (greater than 0.99 in this case)? There are different ways to provide a point estimate of centrality from the posterior distribution such as mean, median and mode (or maximum a posteriori probability (MAP)) under the Bayesian framework, while such point estimates are usually the same due to symmetry under the theoretical distribution of NHST. However, Bayesian inferences tend to emphasize more the uncertainty of an effect, not its point estimate. The green and yellow tails of the posterior density mark the extent of statistical evidence associated with the two specific (two-sided 95% and 90%) uncertainty intervals, and the dotted dark green line shows the median or 50% quantile of the posterior density. Notice 1) the *x*-axis is different between the two densities (standardized value for NHST and effect in physical dimension for Bayesian paradigm), 2) *P* (data | zero effect) under NHST is conceptually and numerically most of the time not the same as 1 − *P* (effect *>* or *<* 0 | data) under the Bayesian framework, 3) the null distribution under NHST has a smooth and regular shape due to the assumption of a standard curve in the model as a prior while the irregular posterior distribution is formulated through random samples through Markov chain Monte Carlo simulations under Bayesian framework, and 4) compared to the conventional confidence interval that is flat and inconvenient to interpret, the posterior density provides much richer information such as spread, shape and skewness.

The Bayesian framework aims to answer a different, though, related question: What is the probability of a research hypothesis *H*_*R*_ based on the observed data, *P* (*H*_*R*_ | data)? Note the difference in what is being “measured” and what is “given” in this proposition, as opposed to the preceding NHST case. Such a probability can be computed by using Bayes’ rule (Kruschke, 2010). In a typical setting the research hypothesis *H*_*R*_ refers to an effect or parameter *θ* (e.g., “population mean”) being positive or negative (e.g., *H*_*R*_ : *θ >* 0). An attractive property of this framework is that it is not typically formulated to generate a binary decision (“real effect” vs. “noise”, or “significant” vs. “not significant”) but instead to obtain the entire probability density distribution associated with *P* (*θ* | data) (Fig. 5, right). This posterior distribution is interpreted in a natural way even though it may take getting used to for those who are unfamiliar with Bayesian inferences. For example, *P* (0.1 *< θ <* 0.2 | data) is the area under the curve within the effect interval (0.1, 0.2). For convenience, one can also indicate “tail probabilities” associated with values commonly used in the literature. For example, the two-sided area indicated in green is 0.05 and the two-sided area indicated in yellow-plus-green is 0.1; these would be analogous to 95% and 90% confidence intervals, respectively, in the NHST framework. Here, we see that the value *θ* = 0 lies inside the left green tail; thus, the probability that the parameter exceeds zero is *P* (*θ >* 0 | data) = 0.99. One can then use the probability in question to *emphasize* or summarize the extent (e.g., “strong”, “moderate”, “weak”, or “little”) of statistical evidence. However, we stress that the goal is to quantify and qualify the evidence, not to make a binary decision in terms of “passes threshold” versus “fails to pass threshold”. In this manner, the value of *P* (*H*_*R*_ | data) is not used for declaring that a result is “real” based on a threshold, but as the amount of evidence on a continuum. There is no need for thresholds and indeed one is encouraged not to use them – in the end they are arbitrary. Under the conventional paradigm, most software implementations do not reveal to the analyst the results that fail to survive the hard threshold (e.g., 0.05). In contrast, we propose a more fluid approach to categorizing statistical evidence: (1) one does not have to adopt a rigid threshold, especially when supporting information exists in the literature; (2) reporting the full results helps portray the entire spectrum of evidence.

An additional benefit of the Bayesian framework is that it provides direct interpretations of statistical evidence on the effects themselves. To see the difference between this and the NHST case, consider the representative distributions in Fig. 5. The left distribution shows the case of a common NHST statistical inference, such as a Student’s *t* under the null hypothesis and the statistic value (shown as a gray box). Note that the abscissa of unitless “t” values is unrelated to physical measures of the parameter. Furthermore, the distribution shape is entirely independent of the data and effect estimates except for the number of degrees of freedom (DFs); any sample with 20 DFs would have the same *t* distribution, regardless of the data and model fit. The inference hinges on the statistic value (gray box) relative to the null distribution. These properties can be compared and contrasted with the statistical inference under the Bayesian framework (Fig. 5, right). Here, the focus is directly on the effect (e.g., activation) in its original scale with a non-standard distribution conditioning on data, model and priors.

Within the traditional NHST framework, one can consider confidence intervals. The use of confidence intervals, however, tends to be plagued by conceptual misunderstanding and even more experienced researchers appear to struggle with its proper interpretation (Morey et al., 2016). For example, one may easily confuse a parameter with its estimator under the conventional statistical framework: a parameter (e.g., population mean) is considered a constant or a fixed effect (without uncertainty) while its estimator (e.g., sample mean) constructed with sampling data is treated as a random variable (with uncertainty). One can assign a distribution to the estimator, but one cannot do so for the corresponding parameter, even if a one-to-one correspondence can be established between the confidence interval of the parameter and an acceptance/rejection region for an estimator of the parameter; this is an all-too-common misconception under NHST. In comparison, all parameters are considered random under the Bayesian framework, and the quantile intervals for a parameter are more directly associated with the corresponding posterior density.

### Limitations of NHST applications in neuroimaging

Much has been written about NHST over the past few decades. Here, we briefly enumerate a few aspects that are relevant in the present context.

#### (1) Vulnerability to misconception

Defined as *P* (data | *H*_0_), the *p*-value under NHST measures the extent of “surprise” under the assumption of null effect *H*_0_ (Fig. 5, left). In contrast, an investigator is likely to be more interested in a different measure, *P* (*H*_*R*_ | data), the probability of a research hypothesis *H*_*R*_ (e.g., a positive or negative effect) given the data (Fig. 5, right), which is conceptually different from, but often mistakenly construed as, the *p*-value. The disconnect between the *p*-value and the probability of research interest often leads to conceptual confusion (Nuzzo, 2014).

#### (2) Arbitrariness due to dichotomization

As the underlying physiological or neurological effect is in all likelihood intrinsically continuous, the introduction of arbitrary demarcation through hard thresholding results in both information loss and distortion. Due to the common practice of NHST and the adoption of significance level as a publication-filtering criterion, a statistically non-significant result is often misinterpreted as a non-existent effect (the absence of evidence is equated with the evidence of absence). The typical implementations in neuroimaging would not allow the user to have the chance to visualize any clusters that are deemed to be below the preset threshold per the currently adopted correction methods; in addition, false negative errors under NHST are largely ignored in the whole decision process regardless of auxiliary information such as the literature and homologous regions between the two hemispheres. Indeed, although even fairly introductory students of statistics will explicitly be aware of this problem, in experimental literature, the practice of describing non-significant results as non-existent effects is puzzlingly widespread.

Hard thresholding leads to conceptual issues such as the following one: Is the difference between a statistically significant result and a non-significant one itself statistically significant (Fig. 6)? Translated into the neuroimaging context, when blobs of spatial units are relatively near but below the adopted threshold (given the cluster size or the integration of cluster size and statistical evidence), available software implementations hide them from the investigator (they are left uncolored, much like a location inside a ventricle). Should those “activation blobs” really be considered as totally devoid of evidence?

**Figure 6:**
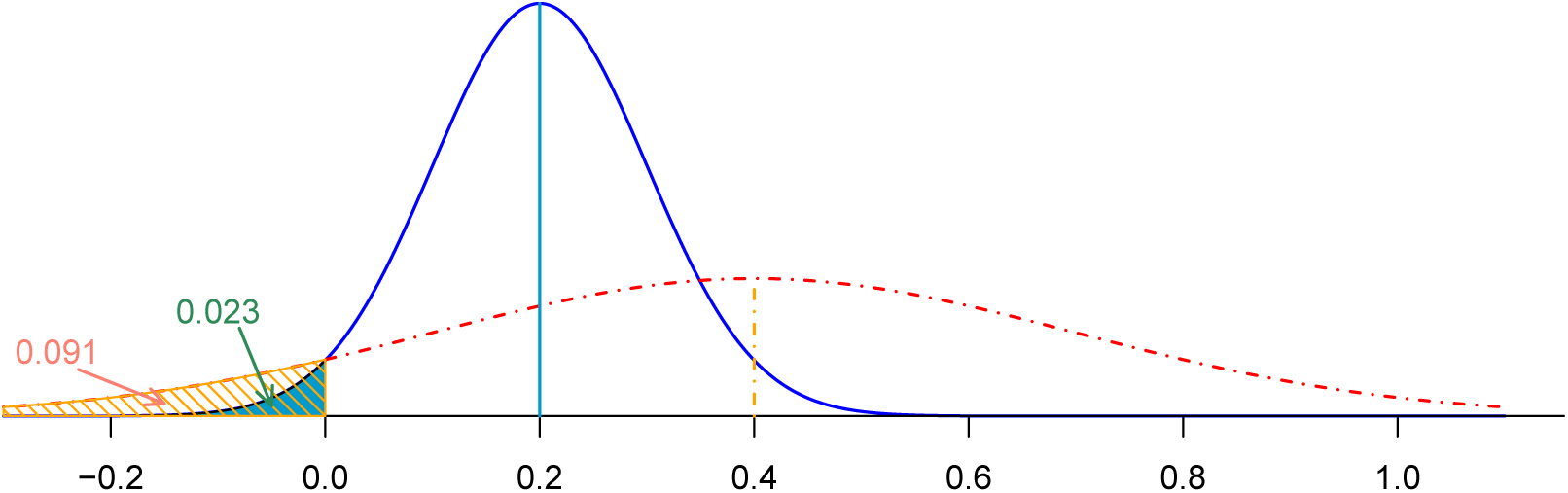
Arbitrariness resulting from dichotomization. Two hypothetical effects that independently follow Gaussian distributions: 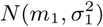 (blue) and 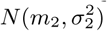 (red), respectively, where *m*_1_ = 0.2, *σ*_1_ = 0.1, *m*_2_ = 0.4, *σ*_2_ = 0.3. Under NHST the first effect would be considered statistically significant with a one-sided (or two-sided) *p*-value of 0.023 (or 0.045); in contrast, the second effect would not be viewed statistically significant given a one-sided (or two-sided) *p*-value of 0.091 (or 0.18). On the other hand, the difference between the two effects is not statistically significant with a one-sided *p*-value of 0.26; in fact, the second effect is more likely larger than the first one with a probability of 0.74.

#### (3) Arbitrariness due to model space

When the whole-brain voxel-wise analysis fails to reveal the expected regions at currently acceptable levels of statistical evidence, how should the experimenter proceed? Indeed, investigators may feel that it is justifiable to adopt “small volume correction” by limiting the data domain, somehow; for example, by focusing on regions of interest outlined in prior literature, or possibly particular regions of outstanding interest (such as the nucleus accumbens in the domain of reward processing). But how to proceed in those circumstances is unclear as, in reality, few researchers are prepared to publicly describe their work as “purely exploratory” at that point. Essentially, the vulnerability to model space manipulations is a byproduct from the current adjustment criterion adopted for handling multiplicity.

#### (4) Overstated estimates and distorted inference

Reporting only statistically significant results tends to produce overestimated or inflated effect sizes (Cremers et al., 2017), as illustrated in the kidney cancer example (Fig. 3); in fact, the current publication practice of only allowing for results with stringent statistical evidence leads to many problems (Amrhein et al., 2019). Suppose that a brain region cannot survive the currently accepted correction criterion (cluster- or permutation-based) but still presents moderate amount of statistical evidence. Such result would have difficulty in getting reported under the current reviewing process, and its absence from the literature may waste the important information that could be utilized as evidence for future studies. If the region fails to reach the designated statistical criterion among four out of ten such studies, the impact of dichotomization on meta analysis could also be substantial. We believe that the reporting of results and publication acceptance should not be based on a dichotomous decision rule in terms of a predetermined statistical significance level (e.g., 0.05).

#### (5) Disregard for effect size and unavailability of uncertainty measures

Because neuroimaging studies perform inferences across tens of thousands of spatial locations, the practice in the field is to present maps that employ colors symbolizing statistical values (Chen et al., 2017). The unavailability of effect sizes leads to crude meta analyses, and renders power analysis challenging in neuroimaging. Furthermore, there is a contradiction, disconnection or inconsistency with the reported results under the conventional analysis approach in the literature. On one hand, due to the unbiasedness property, the effect estimate is considered trustworthy while the corresponding statistic (e.g., *t*-value) and the associated uncertainty (standard error or confidence interval) are frequently not directly interpretable at the spatial unit level because of the extra step required to adjust for multiple testing. On the other hand, due to this lack of interpretability, one cannot take at face value the reported maps with color-coded statistical evidence or the cluster tables with statistical values at peak voxels. In fact, each spatial clique goes through a binary process of either passing or failing the surviving criterion (e.g., either below or above the FPR of 0.05); accordingly, no specific uncertainty can be assigned to the effect at the spatial unit (or peak voxel) level.

#### (6) Lack of spatial specificity

Whereas the conventional massively univariate approach attempts to determine statistical evidence at each spatial unit, the procedures to handle multiple testing render statistical inference viable only at the clusters level. Surviving clusters have different spatial extents, and in practice one may observe some that are quite large. As the unit of statistical inference is the entire cluster, the investigator loses the ability to refer to individual voxels, as shown in the common practice of locating a region via its “peak” voxel. In other words, when a cluster spans a few anatomical regions, spatial specificity may be compromised as the investigator typically identifies only one region in which the “peak” voxel resides.

#### (7) Penalizing intrinsically small regions

Compared to the excessive Bonferroni correction, neighborhood-leveraging procedures offer a relatively effective approach at handling multiplicity. However, the conventional massively univariate approach inefficiently models each individual spatial unit separately, and the compensation for the incurred multiplicity cannot fully recover the lost efficiency, resulting in loss of statistical power as the process attempts to achieve nominal FPR levels. The combination of statistical evidence and spatial extent adopted in recent permutation-based methods (Smith and Nichols, 2009; Cox, 2019) provides a principled approach to address the arbitrariness of primary thresholding, but the approach still discriminates against spatially small regions. For example, between two brain regions with comparable statistical strength, the anatomically larger one would be more likely to survive; and between a scenario involving an isolated region and another with two or more contiguous regions, the former may fail to survive the current filtering methods even when locally exhibiting stronger statistical strength.

Variations of FDR correction have been developed over the years to handle spatial relatedness in the brain (e.g., Leek and Storey, 2008). However, FDR as an adjustment method for multiplicity shares the same issues as the FWE approach (noted above). For example, when effects are not likely to be truly zero, or when the distinction between zero and non-zero is blurry, the FDR control of “false discovery” (zero effect) shares the same logic as the NHST framework, which is based on the idea of a binary distinction between “true” and “false” effects (instead of graded effect magnitudes). Additionally, it is also a separate criterion applied to “fix” the overall results, rather than a coherent strategy, such as the single BML model. Therefore, most of the issues enumerated above apply to FDR correction methods, since FDR ultimately results in adjusted p-values which are then thresholded.

## Applications of Bayesian multilevel modeling in neuroimaging

We now illustrate the applications of Bayesian multilevel modeling to neuroimaging data. We perform group analysis as a prediction process for all spatial units simultaneously in a single model, which assesses the statistical evidence for the research hypothesis *H*_*R*_ based on the available data, that is, *P* (*H*_*R*_ | data).

### Region-based analysis

In a recent study (Chen et al., 2019b), we developed a BML framework for region-based group analysis. As BOLD responses approximately share the same scale, the approach allows information to be pooled and calibrated across regions to jointly make inferences at individual regions. The approach was applied to an FMRI dataset of 124 subjects, each of which had effect estimates at 21 regions. For each individual, a seed-based correlation analysis was performed with the right temporo-parietal junction as the seed (i.e., the correlation between each region and the seed). The question of interest was as follows: What is the relationship between such correlations and individual differences in a questionnaire-measure of theory of mind?

For comparison, the same data were analyzed via conventional whole-brain voxel-wise analysis, which illustrated the difficulty of detecting surviving clusters. With a primary voxel-wise *p*-value threshold of 0.05, 0.01 or 0.005, four clusters were observed after correction through Monte Carlo simulations (Cox et al., 2017); with a primary threshold of 0.001, only two clusters survived (Xiao et al., 2019). In contrast, with BML analysis 8 out of the 21 regions exhibited considerable evidence (Fig. 7). In a Bayesian framework, effect inference is summarized via its entire distribution, the so-called posterior density distribution. (Note that as there is no closed-form solution for the distribution, it must be determined via computational algorithms, such as Markov chain Monte Carlo (MCMC) sampling methods.) Similar to traditional approaches, one can highlight distributional tail areas to denote quantiles of interest (such as 95%), which one may use to label effects as “moderate”, “strong”, and so on. Consider the case of the R TPJp region (top, middle panel of Fig. 7). In this case, the probability of the effect being less than or equal to zero (blue vertical line) is rather small, as it is clearly within the green (0.05) tail. In other words, the probability that the effect is greater than 0 is the area to the right of the blue line (in this case, 0.99). In contrast, in the case of the R Insula (top, right panel of Fig. 7), the probability that the effect lies in the central white area of the plot (which includes zero) is 90%. Now consider the case of the L SFG region (second row, middle panel of Fig. 7). One possibility is to disregard this region given that the zero-effect line does not reside in the colored tails, which is the approach typically adopted under the common practice of dichotomizing. But close inspection of the posterior reveals that the probability that its effect is greater than zero is about 0.93, and the experimentalist may thus wish to deem the results as providing a “moderate” amount of statistical evidence. Regardless of the linguistic terms utilized, one does not have to dichotomize the results, and in fact the investigator is encouraged to report the full posterior^7^. It is also instructive to explicitly compare BML results with those obtained with conventional GLM estimates (Fig. 8). We can see that the BML estimates are pulled to the center (with the exception of region R TPJp), an overall behavior analogous to the one described for the basketball shooting performances.

**Figure 7:**
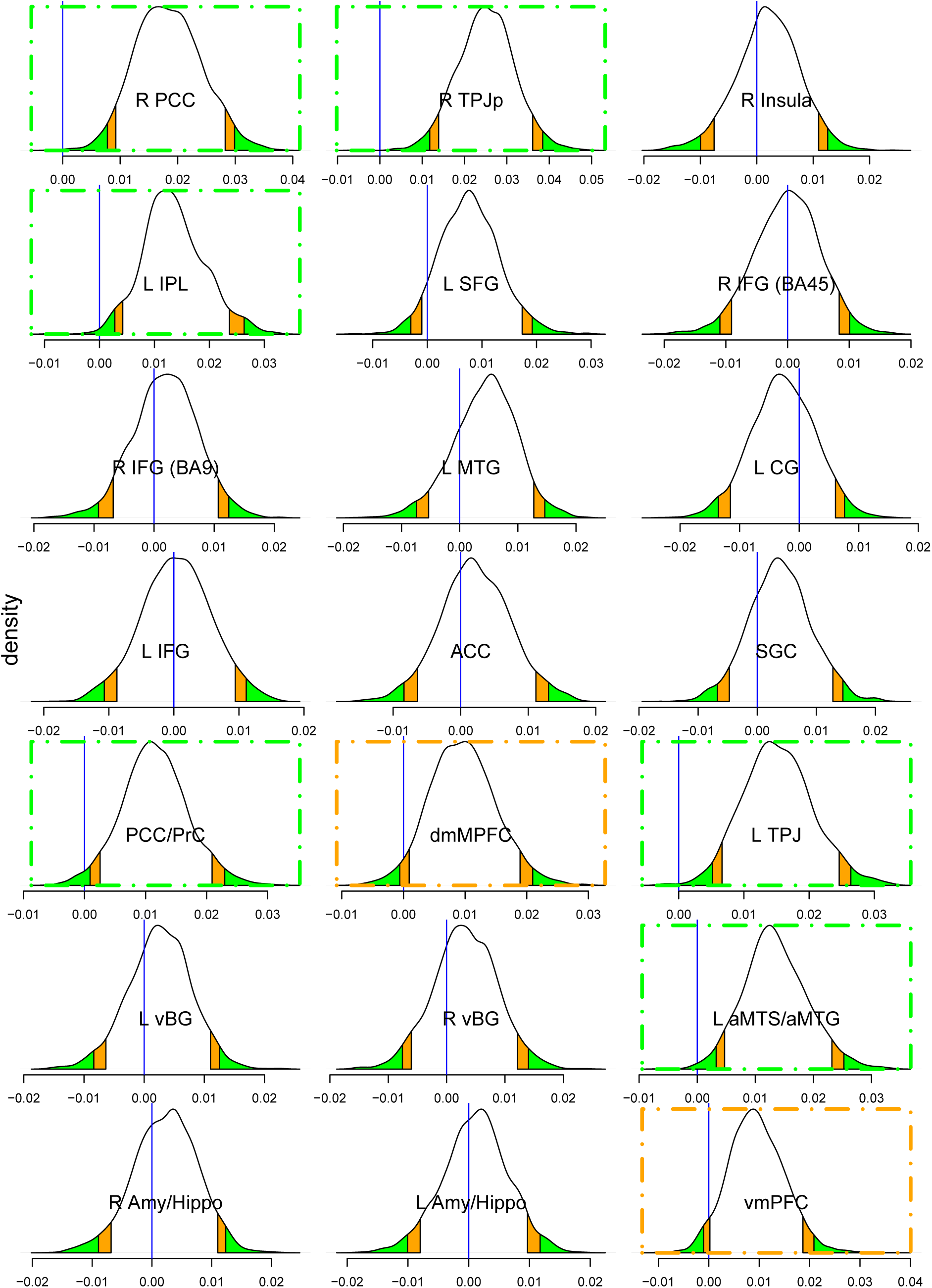
Posterior distributions of the slope effect derived from BML with 124 subjects at 21 regions. The density plot and the associated posterior interval at each region were based on random draws from the same overall high-dimensional posterior distribution that was numerically simulated from the BML model. The vertical blue line indicates zero effect; orange and green tails mark the regions beyond the 90% and 95% uncertainty (compatibility or quantile) intervals, respectively. If results highlighting is desirable, one can claim the regions with strong evidence of slope effect as the blue line being within the color tails, as indicated with orange and green dot-dashed boxes. Compared to the conventional confidence interval that is flat and inconvenient to interpret, the posterior density provides much richer information about each effect such as spread, shape and skewness. Relative to the conventional whole-brain voxel-wise analysis that rendered with only two surviving clusters (Xiao et al., 2019) based on the primary voxel-wise *p*-value threshold of 0.001, the BML showed a much higher inference efficiency with 8 regions that could be highlighted with strong evidence. To illustrate the conventional dichotomization pitfall through a common practice of thresholding at 0.05, the region of L SFG also elicited some extent of slope effect with a moderate amount of statistical evidence: the probability that its effect is greater than zero is about 0.93 conditional on the data and the BML model. Reprinted from Chen et al. (2019b).

**Figure 8:**
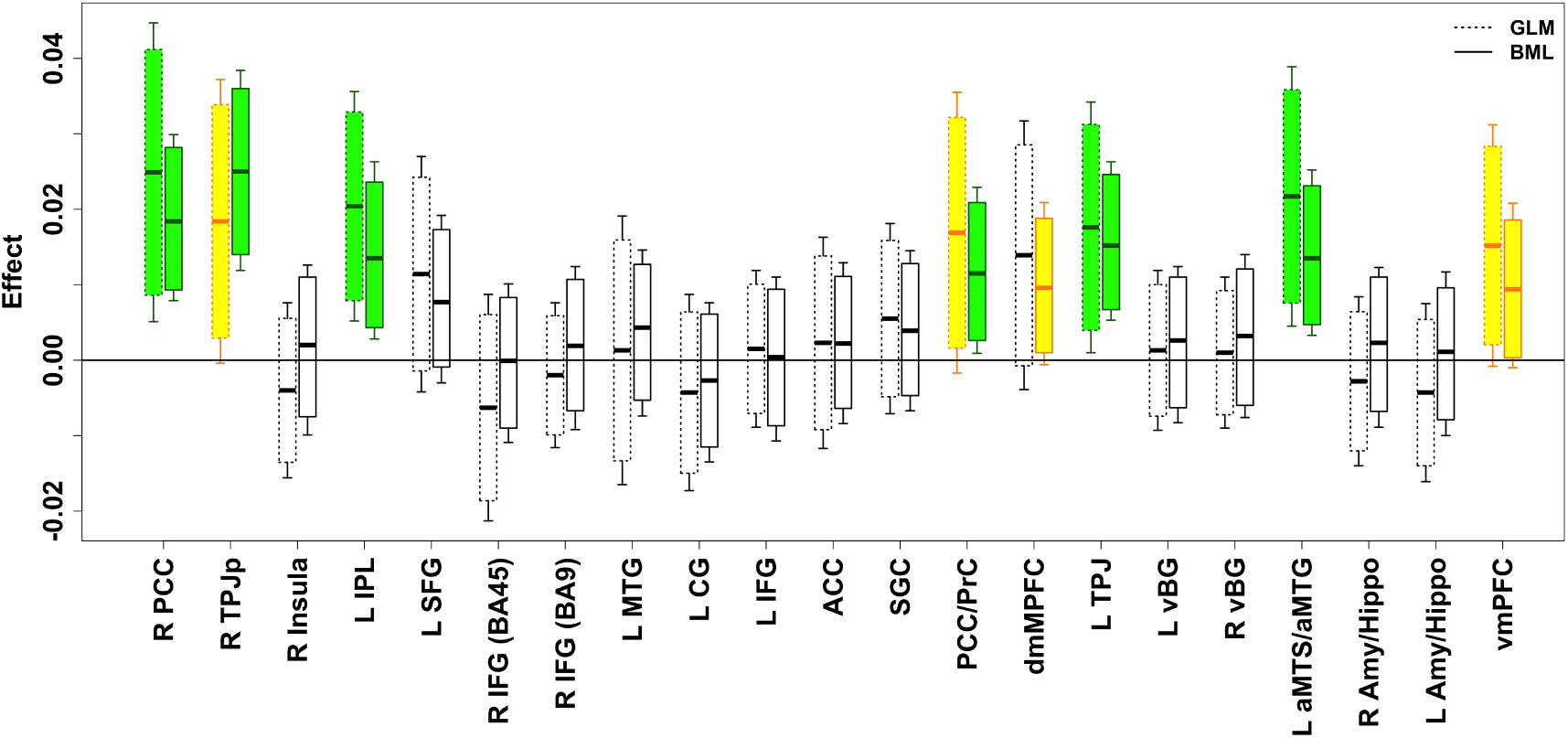
Comparisons of results between the conventional region-based GLM and BML in box plots. For each region, the left bar shows the GLM result that was inferred with the region as an isolated entity while the right bar corresponds to the BML inference. With one GLM for each of the 21 regions, none of the regions would survive FPR correction under NHST. The slope effect is shown with the horizontal black bar in the middle of each box as the mean for GLM or median for BML; each box (or whisker pair) represents the 90% (or 95%) confidence interval for GLM (dashed, left) or uncertainty interval for BML (solid, right). The shrinkage or pooling impact of BML calibrates and “drags” most, but not all regions (see R TPJp). Reprinted from Chen et al. (2019b).

An appealing aspect of Bayesian modeling is that the performance and efficiency of a model can be readily compared to alternative models with the use of cross validation tools such as information criteria and posterior predictive checks. The intuition of visual examination on predictive accuracy is that, if a model is reasonable, the data generated based on the model should look similar to the raw data at hand. For example, the massively univariate approach with GLM yielded a poor fit (Fig. 9A), whereas a much better fit was obtained with BML (Fig. 9B).

**Figure 9:**
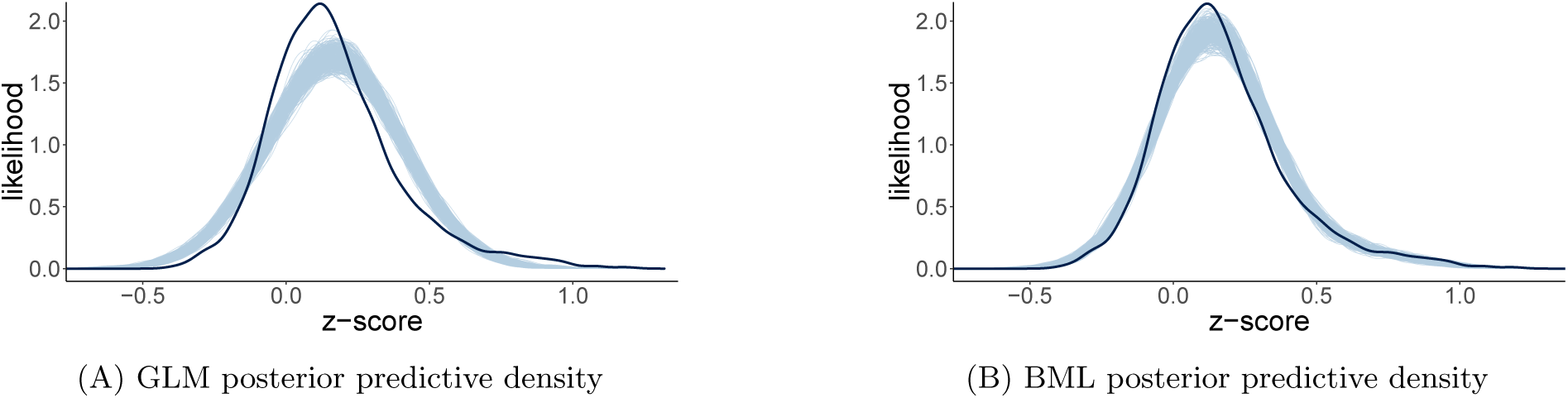
Model performance comparisons through posterior predictive checks and cross validations between conventional univariate GLM (A) and BML (B). The subfigures A and B show the posterior predictive density overlaid with the raw data from the 124 subjects at the 21 ROIs for GLM and BML, respectively: solid black curve is the raw data at the 21 ROIs with linear interpolation while the fat curve in light blue is composed of 500 sub-curves each of which corresponds to one draw from the posterior distribution based on the respective model. The differences between the solid black and light blue curves indicate how well the respective model fits the raw data. BML fitted the data clearly better than GLM at the peak and both tails as well as the skewness because pooling the data from both ends toward the center through shrinkage clearly validates our adoption of BML. To make performance comparisons possible, the conventional univariate GLM was Bayesianized with a noninformative prior (i.e., uniform distribution on (−∞, + ∞)) for the regions. Reprinted from Chen et al. (2019b).

### Matrix-based analysis

We now illustrate the BML framework in the context of another type of data that are of great interest to neuroimaging investigators, namely those summarized in *matrix* form. The most common example is that involving time series correlations between regions (e.g., as in resting state studies), inter-subject correlations among subjects (e.g., as employed for naturalistic scanning (Chen et al., 2019d)), or similarity measures (e.g., representational similarity, latent semantic analysis). In a typical case, an investigator may have tens or even hundreds of ROIs and wish to examine the correlation structure in the data. Consider the case in which the brain is partitioned into 100 ROIs, leading to a 100 × 100 correlation matrix. Given symmetry, one is interested in 100 ×99*/*2 entries (or more generally *n*(*n*−1)*/*2 with *n* ROIs). Although the number of ROIs here is not particularly high, the number of pairwise inferences is 4950, which entails a non-trivial correction for multiple testing. Along the lines described in the preceding section, we developed an approach to model all pairwise correlations simultaneously. Given that a single model is involved, correcting for multiplicity is not required.

The central issue of matrix-based data analysis is to make inferences for each *region pair* which poses considerable challenges. First, correlation values are, by definition, computed over pairs of random variables (e.g., average BOLD time series for FMRI data), but the existence of “shared regions” implies that some pairwise correlations are not independent, namely they are correlated themselves (Fig. 10A). Because the correlation between regions *R*_*i*_ and *R*_*j*_ and that between regions *R*_*j*_ and *R*_*k*_ share a common region *R*_*j*_, proper modeling requires accounting for such covariance between region pairs. A second challenge concerns of course the problem of multiple testing (Fig. 10B), which has been tackled via permutations, where a null distribution is generated to declare which region pairs survive thresholding (Zalesky et al., 2010).

**Figure 10:**
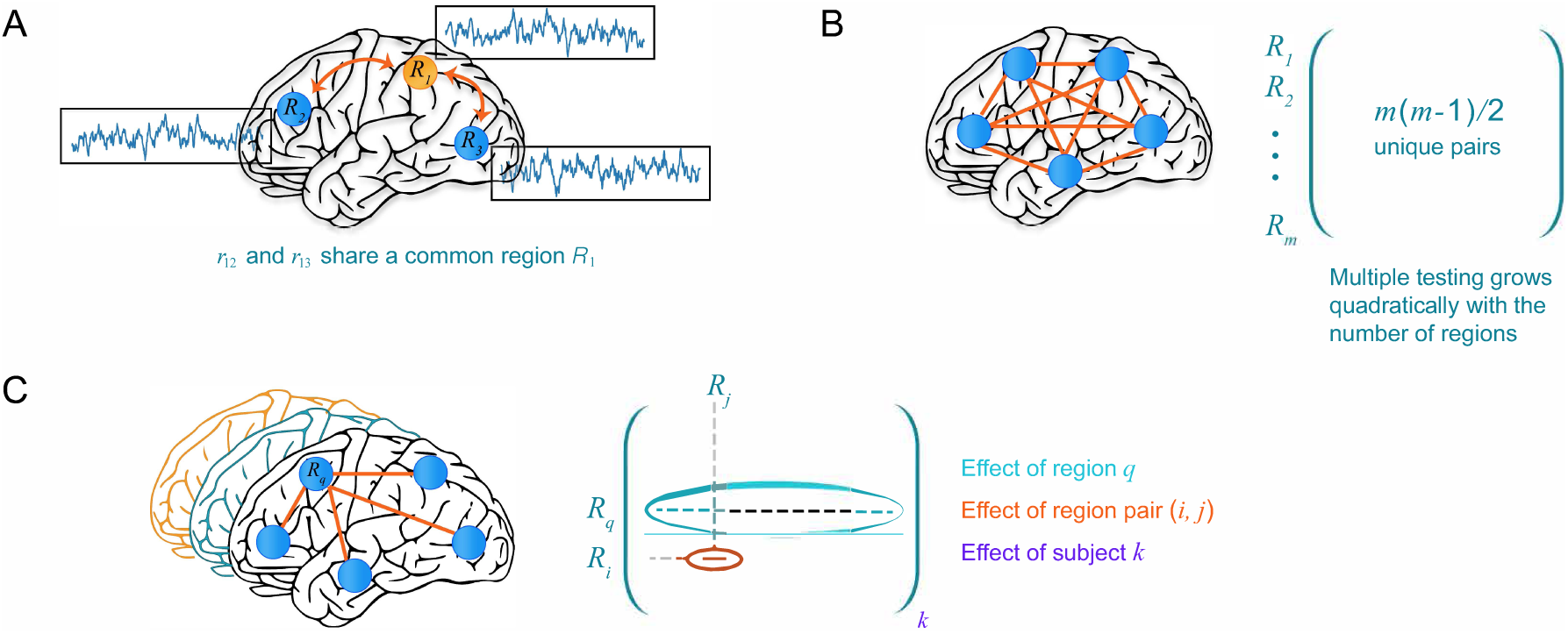
Characterizing the inter-region correlation structure of group brain data. A) Correlations between two pairs of brain regions, *r*_*ij*_, are not independent when they share a common region. Thus, when simultaneously estimating multiple correlations, such relatedness needs to be modeled and accounted for. B) Making inferences about correlations leads to a *multiplicity* problem, in particular how to account for the simultaneous inferences of all effects under NHST. In a Bayesian framework, multiplicity relates to the problem of modeling all correlations simultaneously by invoking information sharing or partial pooling. C) Within a Bayesian multilevel framework, it is possible to frame the problem in terms of capturing the population-level effect of (1) brain region, *R*_*q*_, (2) region pair, *r*_*ij*_, and (3) subject *k*. The characterization of the effect of brain regions is a unique contribution of our framework, which allows investigators to reveal a region’s “importance” within a principled statistical framework. Reprinted from Chen et al. (2019c).

We applied our BML approach to data from a previous cognitive-emotional task (Choi et al., 2012). Briefly, participants performed a response-conflict task (similar to the Stroop task) under safe and threat conditions. A unique feature of the approach developed is that it allows the assessment of effects at the *individual-region level* (Fig. 11). Because the effect of each region pair is decomposed in terms of multiple components including that for each individual region involved, a region-level effect that characterizes the “contribution” of a region can be inferred by BML model. Such an effect, presented with its uncertainty (and the entire posterior distribution), allows the assessment of region importance in a manner that is statistically more informative than those metrics commonly used in graph-theoretic analysis such as degrees and hubs. Although the BML approach is illustrated here in the context of inferences about the population effect for each region and region pair, a researcher might also be interested in group differences, or in the association of a response to a covariate such as one related to individual differences. In the latter situation, one can seamlessly incorporate additional explanatory variables into the BML model, as discussed in the case of region-based analysis in the previous subsection (see also Chen et al., 2019b; Xiao et al., 2019; Chen et al., 2019d).

**Figure 11:**
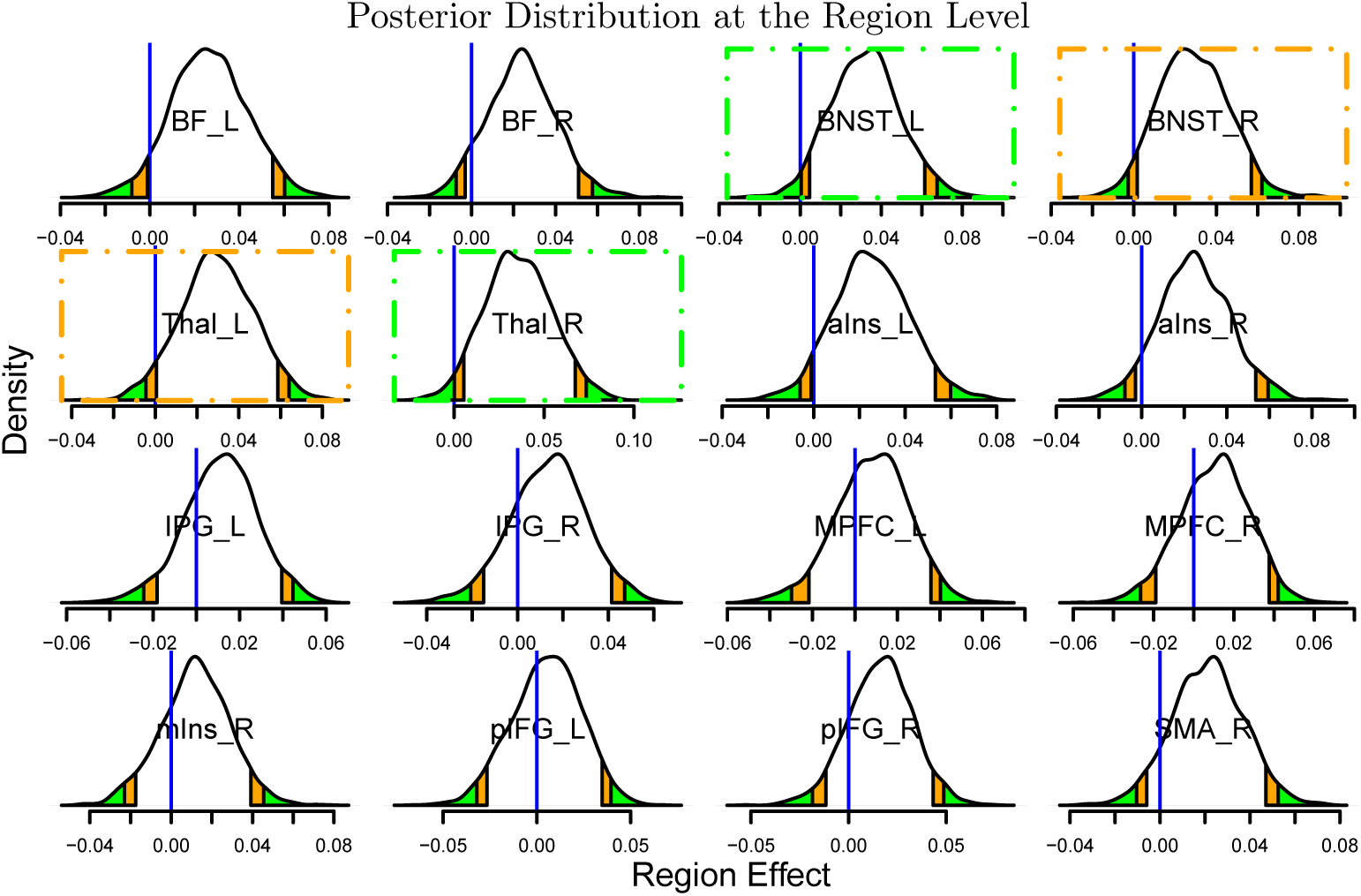
Posterior density plots of region effects (in Fisher’s *z*-value). Each posterior distribution indicates the probability of observing region effects. The orange and green tails mark areas outside the two-sided 90% and 95% quantile intervals, respectively; the blue vertical lines indicate the zero region effect. Consider a region such as the BNST_L (top row, third column): the zero region effect lies in the left green area, indicating that the probability that the effect is positive is greater or equal to 0.975 (conversely, the probability that the effect is negative is ≤ 0.025). The same is true for the Thal_R (second row, second column). In these two cases, there is strong statistical evidence of a region effect, as indicated with green dot-dashed boxes. Two other regions (BNST_R and Thal_L; orange dot-dashed boxes) exhibited moderate statistical evidence of a region effect (the blue vertical lines were each within the orange band). Four more regions forming contralateral pairs of regions (BF_L and BF_R, aIns_L and aIns_R) plus SMA_R also exhibited some statistical evidence as they were close to the typical “convenience” thresholds. Note that the posterior density provides rich information about each effect distribution, including shape, spread and skewness. Unlike the conventional confidence interval that is flat and inconvenient to interpret, it is valid to state that, conditional on the data and model, with probability, say, 95%, the region effect lies in its 95% posterior interval. Inferences about region pairs are shown in Fig. 12. Region abbreviations: BF: basal forebrain; BNST: bed nucleus of the stria terminalis; IFG: inferior frontal gyrus; IPG: inferior parietal gyrus; Ins: insula; MPFC: medial prefrontal cortex; SMA: Supplementary motor area; Thal: thalamus. Other abbreviations: a: anterior; p: posterior; m: medial; L: left; R: right. Reprinted from Chen et al. (2019c).

Figure 12 compares the results based on the conventional GLM approach and those based on our BML proposal. The GLM results were obtained through a massively univariate analysis with (16 × 15)*/*2 independent models, one for each region pair. The BML results were derived by fitting a single model to half of the off-diagonal elements in the correlation matrix. The impact of partial pooling (i.e., information sharing) under BML can be observed by noticing the smaller effect sizes among many region pair. In other words, the effects for most region pairs are “pulled” toward the “middle” relative to their GLM counterparts. The inference efficiency of BML can be appreciated by noticing that 33 region pairs exhibited “moderate” to “strong” statistical evidence; they formed a subset of the 62 region pairs that *individually* were below the 0.05 threshold under GLM. At the same time, when cluster-level correction for multiple testing was applied at FPR of one-sided 0.05 with permutations (Zalesky et al., 2010), no region pairs survived in the GLM case.

**Figure 12:**
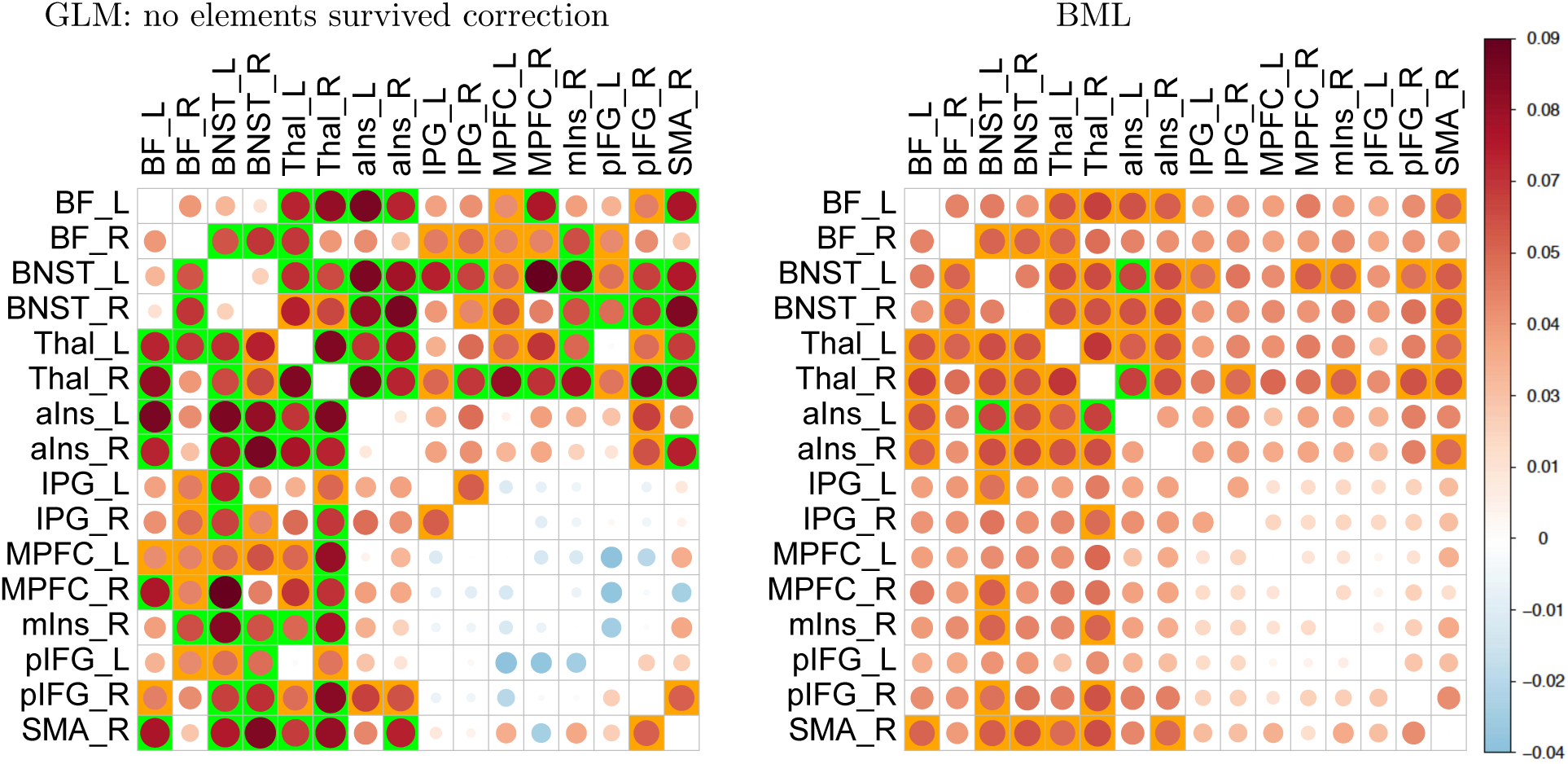
Comparisons of region pair effects between univariate GLM (left) and BML (right) for a matrix dataset. Density plots like Fig. 11 are possible at the region-pair level, but only a summarized version is presented here due to space limitation. The empty entries along the diagonal correspond to the correlation value of 1. The effect magnitude (Fisher’s *z*-value) is symbolized with both circle area and color scheme (colorbar, far right). The impact of partial pooling (or shrinkage) under BML is evident as the effects for most region pairs are “pulled” toward the middle relative to their GLM counterparts. Not to dichotomize but to highlight the extent of the statistical evidence, region pair are colored with a green or orange background based on the strength of statistical evidence associated with 95% or 90% two-sided quantiles. The high efficiency of BML becomes obvious when compared with the conventional GLM. With BLM, 33 region pairs exhibited moderate to strong statistical evidence and they formed a subset of those 62 region pairs declared under GLM. With GLM, 62 region pair were identified as statistically significant (one-sided, 0.05; green and orange boxes) without correction for multiple testing; when cluster-level correction was applied to GLM (FPR of one-sided 0.05 with permutations), no region pairs survived. A distinct feature of BML from GLM, as a natural consequence of the effect decomposition in BML, is the capability of inferring at the region level (Chen et al., 2019c) as presented in Fig. 11. Reprinted from Chen et al. (2019c).

## Discussion

The bulk of statistical training focuses on the conventional NHST paradigm, and so researchers are accustomed to accepting the framework unquestioningly. The “surprise” measure encapsulated by *P* (data | *H*_0_) has become a target goal of data analysis that takes its aim at a “straw man” scenario of “zero effect” under NHST (Fig. 5, left). Unlike situations with truly dichotomous categories (e.g., guilt or innocence in a courtroom trial), most effects or mechanisms under investigation in neuroimaging – and arguably biology more generally – exist along a continuum (e.g., Gonzalez-Castillo et al., 2012), and therefore will be considerably distorted if binarized (as in statistically “significant” vs. “non-significant”).

As discussed by others in the past, a more natural objective of the experimentalist would be to focus on the research hypothesis *H*_*R*_ of interest, which can be assessed through *P* (*H*_*R*_ | data) (Fig. 5, right). With rather benign model assumptions, Bayesian methodology allows just that. Furthermore, advances in Bayesian methods in the past two decades are encouraging a reexamination of some entrenched practices. For example, unbiasedness is generally considered to be a “nice,” if not requisite, property for an estimator. However, it might not always be the most important property. For one, when it is difficult to reach a satisfactory sample size (a nearly universal reality of experimentalists), some extent of statistical bias at the individual-entity level (e.g., participant) might be a cost worth incurring to achieve an overall higher predictive accuracy at the collective level.

Multiplicity is an intrinsic component of the massively univariate approach, and poses substantial challenges, including artificial dichotomy, inflated type M and type S errors, vulnerability to data fishing, suboptimal predictive accuracy and lack of model validation. Moreover, the sole focus on the FPR control under NHST as the rationale of adjusting for multiplicity through neighborhood leverage presumes that the weight of false negatives is essentially negligible in the implicit loss function. These negative consequences are symptomatic of inefficient modeling. To overcome these shortcomings, we propose adopting an integrative BML model and making inferences by sharing information across spatial units. Rather than trying to fight multiplicity through leveraging the relatedness only among neighboring spatial units, we calibrate the information shared across all spatial units through the hierarchical structure of BML: control useful errors of type M and S; improve modeling efficiency; reduce the susceptibility to fishing expeditions; validate each model; and show complete results.

### Modeling frameworks and conceptual differences

An intrinsic distinction exists between random and fixed effects under the conventional statistical framework (e.g., ANOVA and LME). The parameters of interest (e.g., intercept and slopes) under the conventional statistical models (t-test, GLM, ANOVA and LME) are modeled as fixed but unknown effects because these effects are treated as capturing “real” properties of the population (e.g., activation strength, or association between brain activation and behavior). Such effects include the paradigmatic examples with measurement errors for constants such as: players’ performance, epidemiological estimate at the country level, Newton’s *G*, the speed of light *c*, Planck’s constant 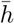, or the effect of interest at each brain region in neuroimaging data. The absence of information about each *fixed effect* is assumed to be temporary, and therefore its uncertainty is considered *epistemic* and will be reduced once the effect can be characterized better and estimated through an improved model (and sufficient data). In this framework, the treatment of fixed effects is “objective” and unbiased since we fully trust the empirical data with the assumption that all potential findings are equiprobable (e.g., activation is a priori as likely to be 0.1% as 100% signal change). In contrast, *random effects* capture intrinsic variations observed in inherently random samples. The inclusion of random effects in the model makes inference generalization possible at the population level, and the uncertainty associated with random-effects variables is treated as aleatoric, reflecting the elusive nature of fluctuations among measuring entities (e.g., subjects). Under the conventional modeling framework, random effects are generally of no interest (e.g., residuals in GLM and subject-specific effects under LME) but are included in the model to account for data variability and to allow generalizations from the current sample to a hypothetical population.

However, such a distinction between random and fixed effects is dissolved under BML by treating *all* effects as random. For example, the effect of interest at each region can be viewed as a varying quantity relative to a collectioin of background information across all regions; that is, by treating the effects among brain regions as the outcome of a “subjective” Gaussian distribution, we no longer need the distinction of fixed versus random effects. Specifically, by leveraging the global assumption of exchangeability among the regions, we infer each region’s effect based on the overall predictive accuracy across all regions. The dissolution between epistemic and aleatoric variability has even been practically applied to effectively modeling measurement errors under the Bayesian framework for conventional physics constants such as the speed of light (Gelman et al., 2014).

To recapitulate, the focus of the Bayesian paradigm is not on the inbuilt metaphysical nature of the effect under study; rather, we treat *all* model parameters (including those that are typically coded as fixed effects under the conventional framework) as random variables by expressing our knowledge and uncertainty through probability distributions. Consider the following example relevant in the context of neuroimaging. For the population of all humans, do we expect there to be a fundamentally fixed, underlying BOLD response in the amygdala (say, 0.538% signal increase) when contrasting images of fearful vs. neutral faces? Should we assume that all the factors contributing to uncertainty of measuring this effect in an experiment (e.g., genetic variability, experimental context, mood, alertness, anxiety, etc.) are epistemic? We suggest that it is reasonable to treat the uncertainty associated with any effect, regardless of the investigator’s interest, as aleatoric within the scope of the BML framework and determine its magnitude uncertainty based on a (large enough) sample of participants. Indeed, the range of aleatoric uncertainty can be reported as a quantile interval from the posterior distribution (or other forms of statistical summary as in conventional approaches). In other words, instead of forcing a view of brain response as exact, intrinsic values across *all* human beings, we make statistical inferences based on an integrative model that renders a representative distribution of the effect. The Bayesian framework thus helps dissolve the distinctions between the uncertainty types. Model parameters such as intercept and slopes are no longer treated as unknown fixed effects at the population level, but are pragmatically considered to be aleatoric, much like random effects such as subjects under the conventional framework. More broadly, a guiding principle of Bayesian statistics is that the state of knowledge about anything unknown should be expressed by a probability distribution.

A little more technically, the move away from assuming epistemic uncertainties (“fixed but unknown”) allows us first to incorporate spatial units as levels of a factor into LME modeling, and then to transition from LME to BML (for technical details, see Chen et al., 2019b). Without operating within the BML framework, the individual spatial units would be typically treated as independent entities (isolated locations) and modeled in parallel under the massively univariate modeling. Consider the example of inferring an effect at each region pair with matrix data. In a traditional LME model each region pair is treated as a random effect, and no inference can be made about individual regions and region pairs other than the intercept (common effect shared by all regions and all subjects), which is typically not very informative in practice. In contrast, within the BML framework, the effect of each region pair is modeled as the contribution of each involved region plus their interaction. In doing so, we can derive both the individual effects *and* their combined contribution as a pair by sampling from the posterior joint distribution (Chen et al., 2019c). Thus, the distinction between fixed- and random-effects characterized in the conventional LME framework is mapped to, in the BML context, the distinction between information pooling across regions and, separately, across-subject variability.

Historically, much has been discussed about the Bayesian approach in the context of “subjectivity” versus “objectivity.” Bayesian methods are frequently linked with the former, and in fact in a negative fashion. As all statistical models are subjective in the sense of idealizing or approximating reality, we favor adopting a pragmatic stance. Certainly, analytic decisions require scientific evaluation, including specific aspects of data processing specifics (amount of spatial smoothing, choice of data included or excluded, model validation, etc.) and uncertainty assignment through a probability distribution. By considering Stein’s paradox and related examples, we hope to have motivated the idea that the “objectivity” of effect inferences (i.e., statistical unbiasedness) should not be the sole criterion in adopting a statistical approach. In fact, the Gaussian prior incorporated in the scenarios of the basketball players, counties, and ROIs, is a model assumption much like those in conventional statistical methods (e.g., Gaussian distribution for residuals and subjects). Note that the Gaussian prior only stipulates the distribution shape, and its specific parameters are actually determined a posteriori through the model conditioning on the data. In contrast, the uniform distribution implicitly assumed under the conventional massively univariate framework, as a special case of BML, is usually not examined or verified in real practice. As experimentalists, we generally have prior knowledge about an effect of interest (e.g., a BOLD percent change of 100% is unrealistic for most contrasts). Adopting a weakly informative prior adds a small amount of real-world information to the model – it provides a “nudge”. Naturally, the analyst should explicitly report all model assumptions, including the priors adopted. Importantly, the performance of a Bayesian model can be evaluated via graphical tools such as posterior predictive checks (Fig. 9) as well as via cross-validation. Taken together, we believe that the global calibration approach can contribute to improving neuroimaging research in multiple ways, including by reducing thresholding-related arbitrariness, increasing model efficiency, and enhancing reproducibility. When BML is applied to the region level, all brain regions are treated on an equal footing based on their respective effect strength. Thus, small regions are not penalized simply because they happen to be small (e.g., less than 20 voxels). In this manner, BML can simultaneously achieve meaningful spatial specificity and detection efficiency. This differs from the conventional analysis, where small regions are inherently placed in a disadvantageous position even if they have similar signal strength as larger ones, as illustrated in Fig. 1.

Related to the current context is the multiplicity issue associated with reproducibility studies as well as with all research studies that employ statistical analysis. For example, one may design a new study to check whether the conclusion from a paper can be duplicated; that is, a replication study intends to determine the generalizability to different subjects, age groups, scanners, sites, preprocessing decisions, analytical tools, etc. One natural question is: under the conventional framework, would one correct for duplicity in statistical inferences? In the same vein, whenever a new statistical analysis is performed, one more inference is added to a huge hypothetical pool of all previous analyses, potentially leading to incrementally inflate the overall statistical evidence: should the analyst incrementally adjust for the significance evidence under the conventional framework through, for example, Bonferroni correction? One may intuitively say no, but how to reconcile the two seemingly contradicting perspectives under NHST? We believe that the Bayesian framework offers a more consistent perspective in this regard. For one particular research conclusion submitted to a journal or reported in the literature, a reviewer or reader may hold a doubtful attitude toward the statistical evidence demonstrated in the paper, reflecting some extent of uncertainty intrinsically embedded in the probabilistic nature of the data and the adopted modeling approach. With more and more similar studies accumulated, we would not expect that all the results are exactly the same; rather, we assume that the studies follow, for example, a Gaussian distribution: most of them are similar to each other with a minority as outliers. In other words, when we emphasize the importance of reproducibility in all scientific investigations, our implicit attitude is homomorphic to embracing, rather than fighting, multiplicity by intuitively applying partial pooling to all the analytical results with a Gaussian prior.

### Potential and limitations of BML applicability in neuroimaging

Table 1 compares the conventional massively-univariate approach and the multilevel framework described here. One noteworthy aspect is that the investigator’s intention concerning the data space has a considerable impact on the extent of statistical evidence in conventional NHST (Kruschke, 2010). For example, if the investigator decides to perform a whole-brain voxel-wise analysis, or to target a few dozen regions of interest, the associated corrections for multiple testing might result in quite different results (from not being able to report a single voxel to reporting noteworthy results across multiple regions). How does BML fare in this respect? Provided that the investigator’s objective is the overall predictive accuracy, partial pooling is expected to produce similar results (Fig. 2D illustrates this behavior for the case of pooling among the top 50 NBA players versus the top 100 players). Accordingly, in a neuroimaging study, if the set of spatial units changes because of an evolving research focus (e.g., adding or subtracting regions), the impact would typically be relatively small as long as the cross-region variability is not substantially smaller than the within-region variability (Chen et al., 2019b). Such adaptivity of the Gaussian prior is supported by ongoing analyses of a task-related dataset with different numbers of regions of interest (e.g., 30, 300, and 1000), resulting in consistent inferences.

**Table 1:** Comparisons of assumptions and properties of massively univariate analysis and Bayesian multilevel modeling

|  | Massively Univariate Analysis | Bayesian Multilevel Modeling |
| --- | --- | --- |
| **Number of models** | number of units plus correction for multiple comparisons | one |
| **Sharing of information** | each unit is independent | units are exchangeable and loosely regularized |
| **Focus of error control** | overall type I (i.e., FPR) | type S (sign) and type M (magnitude) |
| **Strategy for multiplicity** | FPR correction (control for inflated statistical evidence) | partial pooling (control for inflated effect sizes) |
| **Effects uncertainty** | epistemic (effect is intrinsic and fixed, uncertainty from measurement error, etc.) | aleatoric (effect has inherent variability) |
| **Effect inferences** | effect: locally unbiased with no calibration; statistics: uninterpretable at unit level and dichotomized at the clique level | effect: locally biased and globally calibrated; statistics: represented via posterior distribution |
| **Framing of hypotheses** | $P(\text{data} \mid H_0)$: test “surprise” of having the observed data under the null hypothesis $H_0$ scenario | $P(H_R \mid \text{data})$: test evidence for research hypothesis $H_R$ given the observed data |
| **Inference method** | perform NHST with a binary decision based on an FPR-adjusted threshold | assess statistical evidence $P(H_R \mid \text{data})$ through posterior distribution: highlight but no hide |
| **Model efficiency** | local (e.g., unbiasedness of each unit, statistical power) | global (cross-validation, posterior predictive checks) |

We also wish to emphasize the importance of reporting the continuous spectrum of statistical evidence. In the applications described in the previous sections, effect uncertainty was quantified via quantile intervals of the posterior distribution, providing a straightforward and intuitive representation of variability. This approach works well when the number of estimates involved is relatively small (say, 10-20) but is more challenging for larger datasets. Given that neuroimaging data analysis typically involves larger data structures, one may choose to highlight a subset of the effects, particularly those with “strong” or “moderate” evidence. However, these convenient linguistic labels should be operationally defined and not viewed as carrying categorical information (“active” vs. “not active”). There has been a recent call for the distinction between exploratory research (generating research hypotheses with existing data) and confirmatory research (testing existing research hypotheses with new data). Preregistration has been proposed to provide a clearer distinction between the two research aims, and to reduce the influence of publication bias on effect size (Nosek et al., 2018). Our emphasis of complete-results reporting is well-aligned with current trends of study preregistration.

The BML models described here were defined at the level of brain regions. Could the approach be extended and be applied to a whole-brain voxel-wise level? At present the answer is “no” given the computational cost of fitting very large multilevel models. However, this type of analysis may be feasible in the not distant future with the use of Graphical Processing Units and within-chain parallelization (Stan Development Team, 2019). Nevertheless, there are a few disadvantages associated with whole-brain voxel-wise analysis that could be complemented through a region-based approach. For instance, analysis at the voxel level is susceptible to the quality of inter-subject alignment, an increasing challenge as the field keeps pushing for smaller and smaller spatial units (e.g., less than 1 mm isotropic voxels). In addition, inferences based on region-based analysis are region-specific; in contrast, statistical evidence at the voxel level is not necessarily well-aligned with the anatomical or functional definitions, leading to ambiguous or difficult inferences. As region definitions (atlases or parcellations) are available that cover the whole brain (e.g., 1000 parcels (Schaefer et al., 2018)), BML can be directly applied to a set of regions that provide full coverage for the whole brain (Chen et al., 2019d).

The performance of BML requires more testing to assess and validate its consistency and replicability under different scenarios and when applied to multiple datasets. For example, would the inference be consistent when the number of regions increases in real data analysis? Does BML consistently outperform its GLM counterpart in terms of predictive accuracy? The linearity of effect decomposition under BML is a strong assumption, and, as in all linear models, it is an approximation.

In addition, partial pooling under BML is not always effective. On one hand, partial pooling effectively implements a compromise between two “forces,” one that pulls the estimate of the effect toward the center across all regions, and the other toward the local effect at each specific region. Pooling through a weighted average of these two extremes is particularly effective when the across-region variance of the effects is within the same order of magnitude as the within-region variance (sum of cross-subject variance and residual variance). However, when one variance is substantially overwhelmed by the other with a different order of magnitude, then information sharing is basically reduced to one of the two degenerative cases: either “no pooling” (relatively huge within-region variability) or “complete pooling” (relatively negligible within-region variability). Under these scenarios, partial pooling is ineffective, and larger sample sizes are most likely required. For a technical description of these issues, please refer to Chen et al. (2019b).

One potential concern about the BML approach is the underlying assumption about the relationships among the spatial units. Although independence is not required, exchangeability^8^ is needed to guarantee that no differential information among spatial units is available (it also allows approximating the prior distribution across regions in terms of a mixture of identical and independent distributions per de Finetti’s theorem). The assumption of exchangeability is also advantageous because a prior distribution with fewer adjustable parameters will be associated with “sharper” posterior densities (i.e., one with a more discernible peak) than more complex prior distributions (Jefferys and Berger, 1992). Nevertheless, it is to be expected that some regions in the brain will share more information between them (relative to other regions), such as anatomically adjacent or interhemispheric (i.e., homotopic) regions. Therefore, a fruitful direction of research will be to relax the assumption of exchangeability so as to capture potential forms of information sharing between groups of regions. In particular, the multilevel correlation structure among spatial units may be informed by the functional organization of brain regions, such as a gradient between sensorimotor and transmodal areas (Huntenburg et al., 2018). Recent development in the estimation of structured high-dimensional covariance and precision matrices (e.g., Cai et al., 2016) may also help in modeling the hierarchy of spatial relatedness. Another potential modelimg improvement is to adopt a meta-analytic-predictive approach with mixture priors that incorporate historical information through heavy tails (Schmidli et al., 2014). Higher predictive accuracy may be achieved when prior information from the literature can be specifically inserted into the model at the region level.

Another direction for improvement is related to the current practice of splitting the analysis pipeline into at least two levels, one that utilizes one regression model per subject (or even per run/session) to estimate the BOLD signal at the individual level, and another model (e.g., GLM or LME) at the population level. One consequence of such an approach is the potential information loss if the reliability information (e.g., standard error) about the effect estimate is not utilized at the group level. As with other multilevel modeling approaches in the field, BML can potentially take the reliability information into consideration. Thus, a future research direction would be to further integrate the modeling of BOLD data into the BML framework.

Admittedly, there is no magic bullet that could solve all of the problems discussed here. Instead, we may need a cocktail approach that blends multiple solutions. For example, in addition to scientific nuance and preregistration, most likely efficient modeling approaches, model validation, full results and full methods reporting, transparency, and data sharing will contribute to the improvement of reproducibility. We believe that *highlighting* results with stronger evidence without *hiding* weaker ones is healthier than the current sharp thresholding practiced presently. The adoption of BML could also alleviate the vulnerability to data space specification (e.g., what precise set of ROIs is being considered?), as well as promote model validation. In addition, as shown in the kidney cancer example, the focus of controlling for the errors of incorrect sign and incorrect magnitude under BML is potentially more useful than the conventional concept of false positives and negatives under NHST. More generally, a paradigm shift may be needed, one less restricted to null hypothesis testing and more focused on parameter estimation with its associated precision (Gelman and Hill, 2007; Cumming, 2014). Nevertheless, basic statistical principles remain applicable regardless of the modeling approach. For instance, when the across-region variability is substantially smaller than its within-region counterpart, partial pooling through BML may fail dramatically, leading to an undesirable scenario of complete pooling. Per asymptotic theory, such a situation usually indicates a demand for a larger sample size, as may be the case when estimating interaction effects.

## Conclusions

Neuroimaging investigators confront the issue of multiplicity front and center. The analysis challenges posed by this problem are inherently linked to the conventional strategy of modeling the data in a univariate manner: Each spatial unit (e.g., voxel, region) is modelled independently. In the present paper, we described how this approach leads to modeling inefficiency and information loss given the required procedures for handling multiplicity. As an alternative framework, we presented a Bayesian multilevel modeling framework that makes inferences across spatial units by pooling information across them and achieving better overall predictive accuracy. Instead of a multiplicity of models (one for each spatial unit), a single integrative model is employed to embrace multiplicity to modeling advantage. It is our hope that the convergence of theoretical progress combined with algorithm development will stimulate the growth of Bayesian modeling in neuroimaging. Bayesian models also have the potential to encourage reporting experimental findings more fully, instead of dichotomizing them into “significant” and “non-significant” bins, thus reducing information loss in the literature while enhancing both transparency and reproducibility.

## Acknowledgments

The research and writing of the paper were supported (GC, PAT, and RWC) by the NIMH and NINDS Intramural Research Programs (ZICMH002888) of the NIH/HHS, USA. LP’s research is supported in part by the National Institute of Mental Health (R01 MH071589 and R01 MH112517).

## Footnotes

1 The two steps discussed here should not be confused with the “two steps” of first estimating regression coefficients at the individual-participant level and then employing those at a second step of statistical testing across participants.

2 Note that the concept of *true* effect only makes sense under the modeling framework at hand.

3 http://www.fil.ion.ucl.ac.uk/spm/

4 https://fsl.fmrib.ox.ac.uk/fsl

5 Complete pooling can be viewed as specifying a prior with a zero variance.

6 Recall that, in statistics, the bias of an estimator is the difference between this estimator’s expected value and the “true” value of the parameter being estimated. An estimator with zero bias is thus called unbiased, otherwise it is called biased.

7 It is probably advisable to avoid the “codification” of linguistic labels (“moderate”, “strong”, etc.) that are tied to particular probability values (say, 95%). Naturally, considering values along a continuous numerical scale is an advantage of mathematics over discrete language usage. Nevertheless, linguistic descriptors are probably benign as long as the posterior distribution is fully presented.

8 Exchangeability assumes that no differential information is available across the measuring entities (e.g., regions and subjects in neuroimaging) in the model. A set of entities is exchangeable if its joint probability distribution is a symmetric function of the entities; thus, if the order or sequence of the entities is permuted, the joint distribution would not change. Exchangeability captures the distributional symmetry among the entities in a sense that does not require independence: independent and identically distributed set of entities is exchangeable, but not vice versa.

## References

Amrhein, V., Greenland, S., McShane, B., 2019. Scientists rise up against statistical significance. Nature 567, 305–307.

Baggio, H.C., Abos, A., Segura, B., Campabadal, A., Garcia-Diaz, A., Uribe, C., Compta, Y., Marti, M.J., Valldeoriola, F., Junque, C., 2018. Statistical inference in brain graphs using threshold-free network-based statistics. Human Brain Mapping 39(6).

Benjamini, Y., Hochberg, Y., 1995. Controlling the false discovery rate: a practical and powerful approach to multiple testing. Journal of the Royal Statistical Society Series B 57, 289–300.

Bennett, C.M., Baird, A.A., Miller, M.B., Wolford, G.L., 2010. Neural correlates of interspecies perspective taking in the post-mortem Atlantic Salmon: An argument for multiple comparisons correction. Journal of Serendipitous and Unexpected Results.

Bowman, F.D., Caffo, B., Bassett, S.S., Kilts, C., 2008. A Bayesian hierarchical framework for spatial modeling of fMRI data. NeuroImage 39(1):146–156.

Cai, T.T., Ren, Z., Zhou, H.H., 2016. Estimating structured high-dimensional covariance and precision matrices: Optimal rates and adaptive estimation. Electronic Journal of Statistics 10(1):1–59.

Calhoun, V., Adali, T., Pearlson, G., Pekar, J., 2001. A Method for Making Group Inferences From Functional MRI Data Using Independent Component Analysis. Hum.Brain Map. 14:140–151.

Chen, G., Taylor, P.A., Cox, R.W., 2017. Is the statistic value all we should care about in neuroimaging? Neuroimage 147:952–959.

Chen, G., Cox, R.W., Glen, D.R., Rajendra, J.K., Reynolds, R.C., Taylor, P.A., 2019a. A tail of two sides: Artificially doubled false positive rates in neuroimaging due to the sidedness choice with t-tests. Human Brain Mapping 40(3):1037–1043.

Chen G, Xiao Y, Taylor PA, Riggins T, Geng F, Redcay E, 2019b. Handling Multiplicity in Neuroimaging through Bayesian Lenses with Multilevel Modeling. Neuroinformatics (in press). https://rdcu.be/bhhJp

Chen, G., Bürkner, P.-C., Taylor, P.A., Li, Z., Yin, L., Glen, D.R., Kinnison, J., Cox, R.W., Pessoa, L., 2019c. An Integrative Approach to Matrix-Based Analyses in Neuroimaging. Human Brain Mapping (in press). https://doi.org/10.1101/459545

Chen, G., Taylor, P.A., Qu, X., Molfese, P.J., Bandettini, P.A., V, R.W., Finn, E.S., 2019d. Untangling the Relatedness among Correlations, Part III: Extending Model Capabilities of Inter-Subject Correlation Analysis for Naturalistic Scanning. Under revision.

Choi, J. M., Padmala, S., Pessoa, L., 2012. Impact of state anxiety on the interaction between threat monitoring and cognition. Neuroimage 59(2) 1912–1923.

Cox, R.W. 1996. AFNI: software for analysis and visualization of functional magnetic resonance neuroimages, Computers and Biomedical Research, 29: 162–73.

Cox, R.W., Chen, G., Glen, D.R., Reynolds, R.C., Taylor, P.A., 2017. FMRI Clustering in AFNI: False-Positive Rates Redux. Brain Connect. 7(3):152–171.

Cox, R.W., 2019. Equitable Thresholding and Clustering. Brain Connectivity. In press.

Cremers, H.R., Wager, T.D., Yarkoni, T., 2017. The relation between statistical power and inference in fMRI. PLoS ONE 12(11):e0184923

Cumming, G., 2014. The new statistics: why and how. Psychological Science 25(1):7–29.

Derado, G., Bowman, F.D., Kilts, C.D., 2010. Modeling the spatial and temporal dependence in fMRI data. Biometrics 66(3):949–957.

Efron, B., Morris, C., 1976. Families of minimax estimators of the mean of a multivariate normal distribution. The Annals of Statistics 11–21.

Forman, S.D., Cohen, J.D., Fitzgerald, M., Eddy, W.F., Mintun, M.A., Noll, D.C., 1995. Improved assessment of significant activation in functional magnetic resonance imaging (fMRI): use of a cluster-size threshold. Magn Reson Med. 33:636–647.

Gelman, A., Carlin, J.B., Stern, H.S., Dunson, D.B., Vehtari, A., Rubin, D.B., 2014. Bayesian data analysis. Third edition. Chapman & Hall/CRC Press.

Gelman, A., Tuerlinckx, F., 2000. Type S error rates for classical and Bayesian single and multiple comparison procedures. Computational Statistics 15: 373–390.

Gelman, A., Hill, J.L., 2007. Data Analysis Using Regression And MultilevelHierarchical Models. Cambridge University Press.

Glass, G.V., 1976. Primary, secondary, and meta-analysis of research. Educational Researcher. 5(10): 3–8.

Gonzalez-Castillo, J., Saad, Z.S., Handwerker, D.A., Inati, S.J., Brenowitz, N., Bandettini, P.A., 2012. Whole-brain, time-locked activation with simple tasks revealed using massive averaging and model-free analysis. PNAS 109(14):5487–5492.

Huntenburg, J.M., Bazin, P.L., Margulies, D.S., 2018. Large-scale gradients in human cortical organization. Trends in cognitive sciences, 22(1):21–31.

James, W., Stein, C., 1961. Estimation with quadratic loss. Proc. Fourth Berkeley Symposium 1, 361–380.

Jeffreys, W.H., Berger, J.O., 1992. Sharpening Ockham’s Razor on a Bayesian strop. American Scientist 80.

Kang, H., Ombao, H., Linkletter, C., Long, N., Badre, D., 2012. Spatio-spectral mixed-effects model for functional magnetic resonance imaging data. Journal of the American Statistical Association 107(498):568–577.

Kinnison, J., Padmala, S., Choi, J.M., Pessoa, L., 2012. Network analysis reveals increased integration during emotional and motivational processing. J Neurosci. 32(24):8361–8372.

Kriegeskorte, N., Simmons, W.K., Bellgowan, P.S.F., Baker, C.I., 2009. Circular analysis in systems neuroscience: the dangers of double dipping. Nat. Neurosci. 12:535âAS540.

Kruschke, J.K., 2010. Bayesian data analysis. Wiley Interdisciplinary Reviews: Cognitive Science 1(5):658-676.

Leek, J.T., Storey, J.D., 2008. A general framework for multiple testing dependence. PNAS 105(48):18718-18723.

Lindquist, M.A., 2008. The statistical analysis of fMRI data. Statistical science, 23(4):439–464.

McElreath R., 2016, Statistical Rethinking: A Bayesian Course with Examples in R and Stan. Chapman & Hall/CRC Press. https://www.overleaf.com/project/5cb0b178e51a152e6d1cefc1

McShane, B.B., Gal, D., Gelman, A., Robert, C., Tackett, J.L., 2019. Abandon Statistical Significance.

Morey, R.D., Hoekstra, R., Rouder, J.N., Lee, M.D., Wagenmakers, E.-J., 2016. The fallacy of placing confidence in confidence intervals. Psychonomic Bulletin and Review 23(1):103–123.

Nichols, T.E., Holmes, A.P., 2001. Nonparametric permutation tests for functional neuroimaging: a primer with examples. Hum Brain Mapp. 15(1):1–25.

Nosek, B.A., Ebersole, C.R., DeHaven, A.C., Mellor, D.T., 2018. The preregistration revolution. PNAS 115(11):2600–2606.

Pinheiro, J.C., Bates D.M., 2000. Mixed-Effect Models in S and S-plus. Springer-Verlag, New York.

R Core Team, 2019. R: A language and environment for statistical computing. R Foundation for Statistical Computing, Vienna, Austria. https://www.R-project.org/

Santosa, F., Symes, W.W., 1986. Linear inversion of band-limited reflection seismograms. SIAM Journal on Scientific and Statistical Computing. SIAM. 7(4):1307âAS1330.

Schaefer, A., Kong, R., Gordon, E.M., Laumann, T.O., Zuo, X.N., Holmes, A.J., Eickhoff, S.B., Yeo, B.T.T., 2018 Local-Global parcellation of the human cerebral cortex from intrinsic functional connectivity MRI, Cerebral Cortex, 29:3095–3114.

Shaffer, J.P., 1995. Multiple hypothesis testing. Annual Review of Psychology 46:561–584.

Smith, S.M., Nichols, T.E., 2009. Threshold-free cluster enhancement: addressing problems of smoothing, threshold dependence and localisation in cluster inference. Neuroimage. 44(1):83–98.

Stan Development Team, 2019. Stan Modeling Language Users Guide and Reference Manual, Version 2.19.0. http://mc-stan.org

Stein, C., 1956. Inadmissibility of the usual estimator for the mean of a multivariate distribution. Proceedings of the Third Berkeley Symposium on Mathematical Statistics and Probability 1: 197–206.

Stigler, S.M., 1990, A Galtonian perspective on shrinkage estimators. Statistical Science 5(1):147-155.

Tikhonov, A.N., Leonov, A.S., Yagola, A.G., 1998. Nonlinear ill-posed problems. London: Chapman & Hall.

Vehtari, A., Gelman, A., and Gabry, J., 2017. Practical Bayesian model evaluation using leave-one-out cross-validation and WAIC. Statistics and Computing. 27(5):1413–1432.

Whitfield-Gabrieli, S., and Nieto-Castanon, A., 2012. Conn: A functional connectivity toolbox for correlated and anticorrelated brain networks. Brain Connectivity. doi:10.1089/brain.2012.0073

Worsley, K.J., Marrett, S., Neelin, P., Evans, A.C., 1992. A three-dimensional statistical analysis for CBF activation studies in human brain. Journal of Cerebral Blood Flow and Metabolism, 12:900–918.

Xiao, Y., Geng, F., Riggins, T., Chen, G., Redcay, E., 2019. Neural correlates of developing theory of mind competence in early childhood. NeuroImage 184:707–716.

Zalesky, A., Fornito, A., Bullmore, E.T., 2010. Network-based statistic: identifying differences in brain networks. NeuroImage 53:1197–1207.

Zhang, L., Guindani, M., Vannucci, M., 2015. Bayesian models for functional magnetic resonance imaging data analysis. Wiley Interdisciplinary Reviews: Computational Statistics 7(1):21–41.

